# Furrow tillage reduces soil carbon loss and enhances microbial metabolism

**DOI:** 10.1101/2025.08.01.667305

**Authors:** Arnab Majumdar, Munish Kumar Upadhyay, Argha Ghosh, Rakesh Biswas, Ioly Kotta-Loizou, Martin Buck, Mark Tibbett, Biswajit Giri, Debojyoti Moulick, Manoj Kumar Jaiswal, Tarit Roychowdhury

**Affiliations:** Department of Life Science, Faculty of Natural Sciences, Imperial College, London, SW7 2AZ, UK; Centre for Environmental Science & Engineering, Department of Civil Engineering, Indian Institute of Technology Kanpur, Uttar Pradesh-208016, India; Department of Agricultural Meteorology, Odisha University of Agriculture and Technology (OUAT), Bhubaneswar-751003, India; Department of Chemistry, University of Ulsan, 93 Daehak-ro, Nam-gu, Ulsan, South Korea; Department of Sustainable Land Management and Soil Research Centre, School of Agriculture Policy and Development, University of Reading, Reading, RG6 6AR, UK; Department of Earth Sciences, Indian Institute of Science Education and Research (IISER) Kolkata, Mohanpur, West Bengal-741246, India; Department of Environmental Science, University of Kalyani, Nadia, West Bengal, 741235, India; School of Environmental Studies, Jadavpur University, Kolkata, West Bengal-700032, India

**Keywords:** Furrow tillage, global carbon loss, sustainable agronomy, plant physiological changes, microbial molecular response

## Abstract

With increasing global carbon loss from agricultural fields, agricultural practices should be monitored with proper regulatory measures. To address three problems of minimising carbon loss, optimising soil fertility and enhancing soil microbiota together, this study has developed a new innovative “Furrow tillage” approach that affords a balance between conventional and conservative tillage. Based on two years of trials in rice fields at multiple sites, this practice has justifications from agronomy, geochemistry, crop physiology, and molecular microbiology. Furrow tillage illustrated low carbon loss, similar to no-tillage, while retaining high nutrient bioavailability, similar to deep tillage. Microbial diversity, molecular responses and metabolic activities showed that furrow tillage induced microbial interactions while allowing a better entrapment of carbon dioxide in the soil. The study used CO_2_ flux, spatial distribution, total and available elemental analysis, plant ultra-structural observations, microbial metagenomics, high-throughput sequencing and molecular modelling to establish an optimal soil-crop quality, microbial maintenance and reduce carbon loss through furrow tillage practice.

## Main

The primary concern of the modern era is climate change, in which global warming, due to the continuous release of greenhouse gases (GHGs), is a major contributor. The release of CO_2_ from agricultural fields and the carbon use efficiency are interlinked because of diverse agronomic practices where microbes play a critical role in the storage or loss of soil carbon content^1^. Although several tillage methods have been developed alongside established no-till and deep-till practices, the conservation of soil organic carbon (SOC) and its maintenance during intensive farming remains contentious^2,3^. For the no-tillage fields (NTF), soil disturbances are the least compared to the conventional deep-tillage fields (CTF)^4,5^. It is necessary that we understand the limitations and benefits of both these no-tillage and deep- tillage practices to showcase the potential of a new agronomic method that can control CO_2_ release from the field while keeping the soil quality and nutrient dynamics high^6^. Global reviews have described the mechanism of no-tillage farming on SOC, showing that predicted C sequestration in soils was not conclusively found^7–9^. Conventional deep tillage, in contrast, provides a loose tilth for better plant root development and consequent nutrient availability, resulting in greater plant-microbial interactions^5,10,11^, but fails to improve the soil carbon storage. Soil microbial communities are vital in maintaining and cycling this SOC content, which indicates the active metabolism and enzyme turnover in microbial communities^1,12^. It is, however, not clearly shown and explored what might be the effect of differential irrigation practices when combined with the tillage setups. Waterlogging conditions can retain or reduce CO_2_ release, whereas deficit irrigation or alternate wetting and drying (AWD) can trigger CO_2_ release^13^. However, researchers agree that using AWD can benefit the soil by increasing nutrient availability, microbial diversity, metabolism, plant growth responses and reducing heavy metal uptake compared to continuous flooding^14–16^. The aim of this field-based trial was to test these two well-known tillage practices, no-tillage and conventional deep tillage, with the newly developed furrow tillage from agronomic, geochemical, plant physiological, soil microbial and molecular aspects. In this furrow tillage field (FTF), a unique combination of NTF and CTF has been maintained. Therefore, to establish the optimum agronomic practice that can retain SOC stock, increase nutrient availability, promote plant physiology, and increase total microbial community diversity and metabolism, this study proposes to adopt the furrow tillage practice.

## Results

### Carbon variability under differential tillage

Tillage practices certainly modulate the soil carbon pool and the labile fraction of carbon that gets easily altered by the soil properties and microbial communities^17,18^. How the soil carbon pool got altered and released varied CO_2_ content from NTF, CTF, and FTF has been shown in **Fig. 1**. SOC indicates the total changeable carbon pool, where soil chemical properties and microbial activities can produce the labile form of carbon. Under the no-tillage practice, field soil carbon was found to be 10.17-16.07 g kg^-1^ soil, retaining the highest SOC but the lowest amount of labile carbon fraction within the range of 0.22-20.27 g CO_2_ g^-1^ SOC (**Fig. 1a-b**). Following NTF, the moderate SOC retention was associated with the FTF field setup having 7.95-11.04 g kg^-1^ soil with the intermediate labile carbon content of 0.27-0.33 g CO_2_ g^-1^ SOC. The lowest carbon pool was found in the CTF field within a range of 4.44-7.59 g kg^-1^ soil. However, the highest labile form of carbon was also associated with this field, having 0.37-0.42 g CO_2_ g^-1^ SOC. Released CO_2_ from agricultural fields critically depends on the soil-bed disturbances, which transform labile carbon content into CO_2_ due to microbial and geochemical activities^19^. This two-year-long field trial has shown the correlation between the labile form of carbon and the released CO_2_ from the fields. The newly developed FTF practice had a moderate release rate of field-emitted CO_2_ within a range of 3.94-4.38 g C m^-2^ d^-1^, which is closer to the lowest released CO_2_ from the NTF fields within a range of 2.43-2.84 g C m^-2^ d^-1^ (**Fig. 1c**). The highest CO_2_ release was measured from the CTF fields within the range of 6.29-7.11 g C m^-2^ d^-1^. The concentration-based labile carbon and CO_2_ release data were tested for the correlation regression (**Fig. 1d**), which suggests a generally positive trend between these two terms. More soil disturbances in the CTF triggered the geochemical activities having the highest conversion of CO_2_, resulting in the highest R^2^ values in the correlation plot, followed by FTF and NTF.

**Fig. 1.**
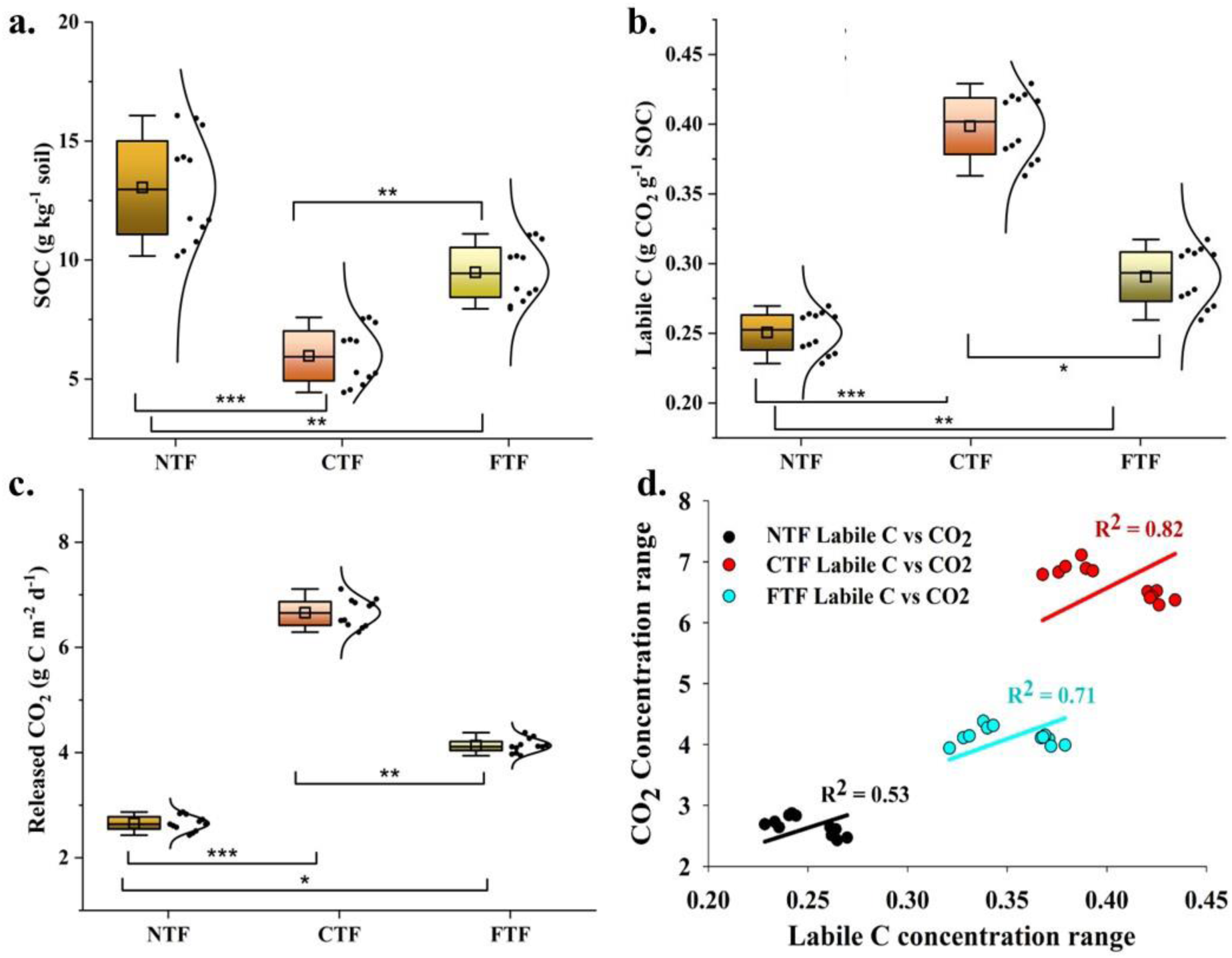
Soil carbon content and released CO_2_ from three differentially tilled fields with a correlative relation. The combined data from flooded and dry-wet field setups are presented here to show the bulk content difference based on the tillage practices. A box-whisker plot has been combined with the distribution curve to indicate the pattern of the data distribution. Total SOC **(a)**, labile carbon content **(b)**, released CO_2_ from the fields **(c)** and regression analysis with linear fit **(d)** of the field-dependent CO_2_ release. Data are representative of the average trend of 2 years of analysis for a triplicate data set. One-way ANOVA with p<0.05 has been performed.

To better understand how much the proposed furrow tillage practice was efficient, the fold change results are shown in the **extended figure 5**. The flooded irrigation had better retention of CO_2_ compared to the AWD practice. However, the margin was low and considering the other benefits of AWD cultivation^20^, this should be practiced more than flooded (FL) cultivation. The fold change of released CO_2_ of 2.44-2.48 from NTF to CTF was the highest, indicating the difference between conventional deep tillage and no-tillage methods. However, the fold change from NTF to FTF showed a 58% decrease compared to the conventional tillage practice, which significantly retains the labile form of the carbon in the soil for microbial activities and nutrient entrapment. The total agricultural field area used for the no-tillage practice is around 507 million hectares, while the practice of tillage is associated with 1150 million hectares, making a significant difference in CO_2_ release, soil quality and crop production^21^. Based on the calculations and projected CO_2_ release from NTF and CTF, the global fold change is shown in the **extended figure 6**. If 30% from the practiced CTF is converted to the FTF, there would be a significant reduction in the global CO_2_ release and soil carbon loss. This projected data trend shows the drastic difference of CO_2_ release from NTF and FTF compared to CTF at the global scale. Pie charts indicate the current and proposed area distribution of tillage practices if FTF is promoted globally. This trend was similar in both years. One-way analysis of variance (ANOVA) was performed for all these datasets to check the statistical significance at the *P*<0.005 level. To get the right understanding and accuracy on data trend, three different post-hoc analyses were performed- Bonferroni correction test, Tukey honestly significant difference (HSD) test and Fisher’s least significant difference (LSD) test. The Origin 2024 version was used, and the **extended figure 7** shows the scores for each of these tests, tested for the inter-group analysis. The figure with red-lined boxes indicates the considered *P*-values at a 0.05 significance level, while scores 0 and 1 indicate non-significant and significant in the mean variance. Results suggested a clear significance in the range and mean values of SOC, labile C and CO_2_ release from NTF to CTF and FTF fields. The average data from both years was tested for the pattern distribution using violin plots in **extended figure 8**. This range of average SOC, labile soil carbon content and released CO_2_ from NTF, CTF and FTF under AWD and FL practices showed the variable positive skewness among these setups. FTF showed a positive, skewed distribution in all three carbon estimations compared to the NTF and CTF, indicating an optimal carbon balance. Points in these violin plots indicate 12 sampling sites. The released CO_2_ data was tested for the standard variance test to uncover the statistical significance of the data using one-way analysis of variance (ANOVA) at a *P*<0.05 level. This set was further considered for two different post-hoc analyses, the Bonferroni correction test and the Mann-Whitney U test. The considered groups for the Bonferroni correction were three, considering TF *vs* CTF, NTF *vs* FTF and CTF *vs* FTF, which provided the corrected Bonferroni α value as 0.0166. All the tested *P*-values were less than this corrected α value, indicating the significance among the three groups. Mann-Whitney test results are also indicative of the strong significance in the inter-group variance with the exact *P*-values and are shown in the **extended tables 2-4**. Lower U values within NTF vs CTF CO_2_ release and NTF vs FTF elemental bioavailability and microbial distribution proved the support to FTF practice over NTF and CTF. The spatial variation of C release in the locality of the experimental fields has been given in **Fig. 2a-b** for two consecutive years 2022 to 2023. The locations of the experimental fields under different tillage practices have also been shown. The thematic layers corresponding to various tillage practices have been represented within a uniform scale ranging from 2 to 10 g C/m^2^/day. The maps clearly indicated a higher degree of C release in the year 2023 across all the tillage practices. These released CO2 maps can be compared to the global carbon emissions from other industrial sectors, other than agricultural practices. This helps to understand the differential pattern of C-release from the regional to the global scale. Thus, the projected values of possible minimised C content from the agricultural fields (as shown in the extended figure 6) can be linked to this global distribution pattern.

**Fig. 2.**
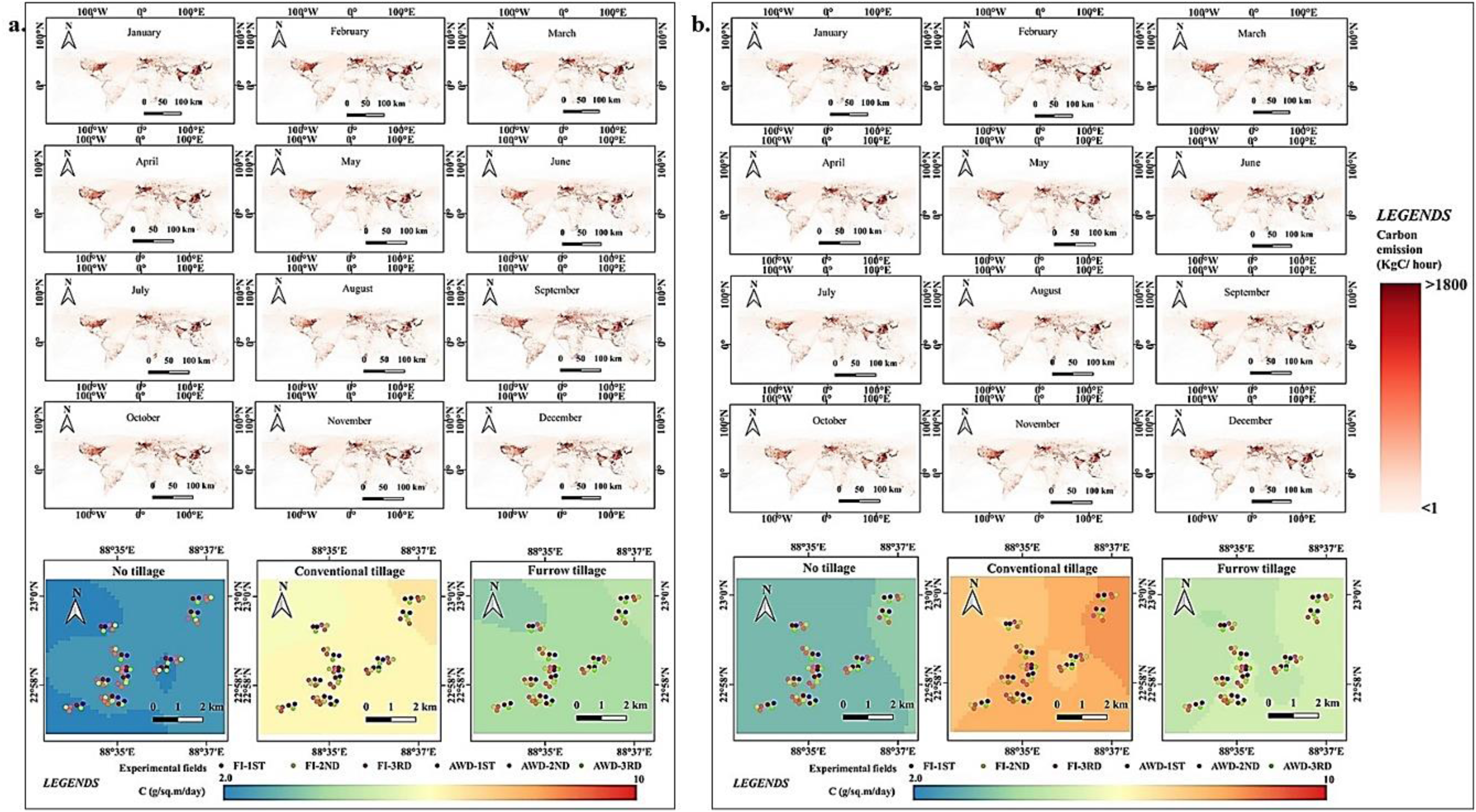
Spatial distribution map of the emitted carbon as per the GRACED database and released CO_2_ in each of the trial fields (no-tillage, conventional deep tillage and furrow tillage) from two consecutive years (**a**-2022; **b**-2023). Colored dots represent three different collection time points (1^st^, 2^nd^, and 3^rd^) during rice cultivation with two different irrigation regimes (flooded_FL and alternate wetting-drying_AWD). Twelve experimental sites are marked here, covering a region where long-term practice of both no-tillage and tillage is known.

Considering the variation in C release with respect to the variation in the tillage practices, no tillage farming recorded the least C release in both years, 2.4 times less than the CTF, followed by furrow tillage farming. Conventional tillage farming resulted in a greater amount of C release. In order to spatially interpolate the C release using the Ordinary Kriging (OK) method^22^, the logarithmic variogram models were selected as the best-fitted model for all the tillage practices during both years. The best-fit variogram model^23^ data for the present study are shown in **extended table 5**. The highest value of the coefficient of determination (R^2^) was obtained from the furrow tillage practices in both 2022 (81.74%) and 2023 (58.41%). No tillage farming demonstrated the least R^2^ values as 53.90% and 35.54% in 2022 and 2023, respectively. The R^2^ obtained in 2023 was found to be lower than that obtained in 2022.

### Stomatal activity and internal anatomy

It is a well-known fact that elevated levels of CO_2_ in the surrounding atmosphere can trigger a plant’s physiology, particularly the vascular structure in the leaf, for a higher passage of water, nutrients, and gaseous exchanges^24^. This study experimentally investigated the effect of released CO_2_ from the field on plant leaf stomatal movement in terms of opening and closing and vascular (xylem and phloem) structural integrity. During the experiment, snap-freezing of the sampled leaf ensured the exact position and status of leaf stomata while another sampling of the same leaf after 2hr incubation helped to compare the changes. **Fig. 3a** shows the stomatal status in NTF, CTF and FTF leaves at 0 hrs, which was considered as the beginning of the experiment, followed by the final status of the stomata from the same leaf in **Fig. 3b** after 2 hrs of incubation. **Extended Figure 9** shows how a CO_2_ molecule enters the plant cell through the stomatal opening when the intra-cellular CO_2_ level is low, and after having an ambient carbon concentration, the guard cells close the stomatal opening. The same leaf for observing such stomatal activity was followed closely in rice plants from all 12 experimental sites during the inflorescence and harvest phases. Changes in stomata activity from **Fig. 1a to 1b** clearly correlate with the degree of available CO_2_ within the surrounding leaves, depending on the field setups. The least CO_2_ was released from the no-tillage practice, resulting in partially opened stomata in the 2 hrs NTF leaf, whereas the CTF leaf stomata were completely closed during this incubation phase. The FTF leaf showed stomatal closure near about 85%, which suggests the release of CO_2_ was enough for the plant’s requirement. The plant’s vascular structures in **Fig. 3c** show the varied integrity within the xylem and phloem formations under NTF, CTF and FTF-grown plants. Studies are evident for the CO_2_ uptake through root xylems^25^ and growing plants at the elevated CO_2_ can increase the xylem-phloem passage^26^. The electron microscopic observation of the vascular structures in rice leaves under the AWD and FL conditions is proven to have altered effects on the xylem, phloem and bundle sheath^27^, indicating the suitability of AWD compared to the FL. This study thus defined the effect of structural features of vasculatures under differential tillage practices. Only AWD-irrigated plants were considered for the structural observation. Yellow arrows point out the xylem and phloem in the vascular pathway, where a mild distortion was observed in the NTF-grown plants. This presumably results from having lower CO_2_ levels within a given time span. Considering the higher accumulation of CO_2_ intracellularly, CTF and FTF anatomical rigidity remained undisturbed.

**Fig. 3.**
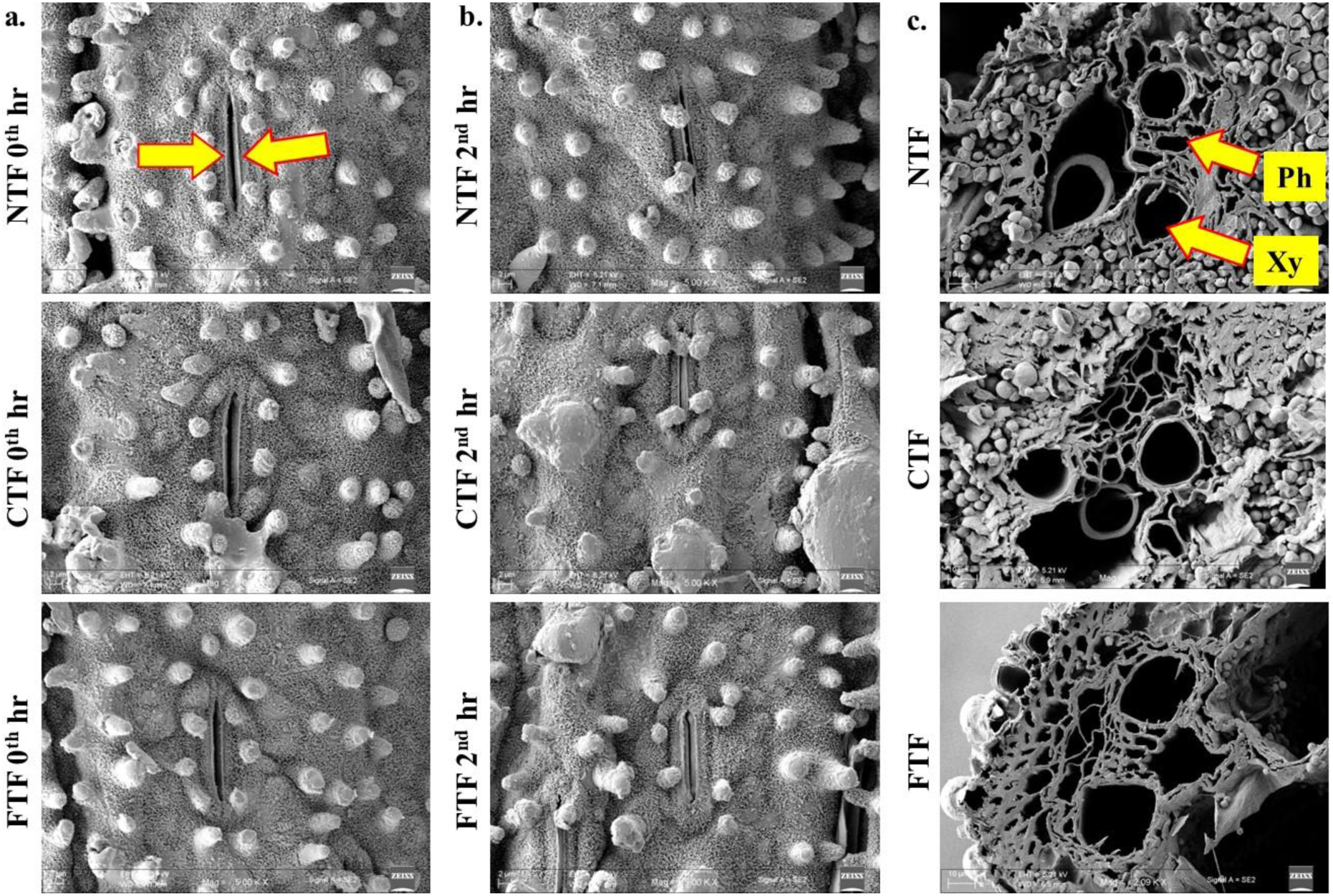
Plant leaf stomatal movement **(a-b)** and its co-establishment with vascular structure and activity **(c)**. Yellow arrows in (a) indicate the guard cells of the stomatal boundary while opened, and in (c) point out the xylem (Xy) and phloem (Ph). Stomatal opening and closure were observed and captured at 2 hr time interval. The sample was collected from the same plant leaf at both times.

### Elemental distribution and bioavailability

While proposing a new agronomic practice, assessment of soil parameters and nutritional values are essential to define, as shown in **extended figure 10** and **Fig. 4**. For rice cultivation, N-P-K, Mg, Ca, and Zn play vital roles^28^ while Cu and S are also important to maintain the healthy soil microbiota in the rice rhizosphere. These selected elements were analysed for their total concentration, and the differences in the contents are presented based on the tillage practices. From the **extended figure 10**, it is clear that AWD irrigation leads to better water retention as compared to FL irrigation. The nutrient mix was relatively higher in CTF and FTF than NTF, except for N, K and Mg. As there was no additional nutrient application other than the regular fertilization by the farmers, it was assumed that the difference in this total elemental content was associated with the soil itself. These concentration changes might be the combined effect of tillage and irrigation regimes. CTF had the highest soil mix, while FTF had some specific mixing along with the furrow channels, which contributed a higher elemental content in the sampled soil from the top 0-20 cm profile. However, NTF was mostly undisturbed with hardened soil clods, and only irrigation regimes had the chance of dispersing partial clods, releasing the elemental content. This hypothesis was further tested by analysing the bioavailable fractions of these three field setups (**Fig. 4**). The sequential extraction method allows the separation of the exchangeable and carbonate-bound fractions from the organic matter-bound, mineral-bound and residual fractions of the soil elemental concentrations. For each of these elements, the highest bioavailability was measured in the CTF soil, followed by the FTF soil. For the NTF, this availability was the least. The violin plot for each element shown in the figure represents the difference in implementing the AWD and FL irrigations combined with NTF, CTF, and FTF. The developed furrow tillage outcome is similar to the CTF soils in terms of high bioavailability but lower surface disturbances. Density cures in these violin plots indicate soil fluctuations with multiple high-mode values. This indicates the heterogeneity of a natural soil sample. The fold changes of bioavailability from NTF to CTF and FTF, as shown in the **extended figure 11**, indicated a higher positive range of fold change in the elemental availability from NTF to CTF. In contrast, the NTF to FTF fold changes were positive but less than the CTF. A negative range of fold change was observed from CTF to FTF. Also, in all three tillage practices, AWD fields were found to have higher elemental availability than the FL fields. The average bioavailability of these selected nutrients in two years showed a positive skewness under FTF field setups compared to the NTF and CTF, as shown in **extended figure 12**. The violin plot showed a normal distribution of these datasets, where a higher distribution range and longer density plot of the violin area were associated with the CTF and FTF compared to the NTF. These data sets suggest the FTF tillage practice combined with AWD irrigation can potentially retain optimal nutrients for plant growth. Considering that plant nutrient uptake largely depends on the bioavailability in the soil, a similar data trend was observed in total plant elemental contents, as shown in the **extended figure 13**. For all the selected elements, NTF-grown rice plants had considerably less nutrients than the FTF and CTF-grown plants. AWD irrigation displayed the advantage of providing higher nutrient availability for uptake than did the FL irrigation.

**Fig. 4.**
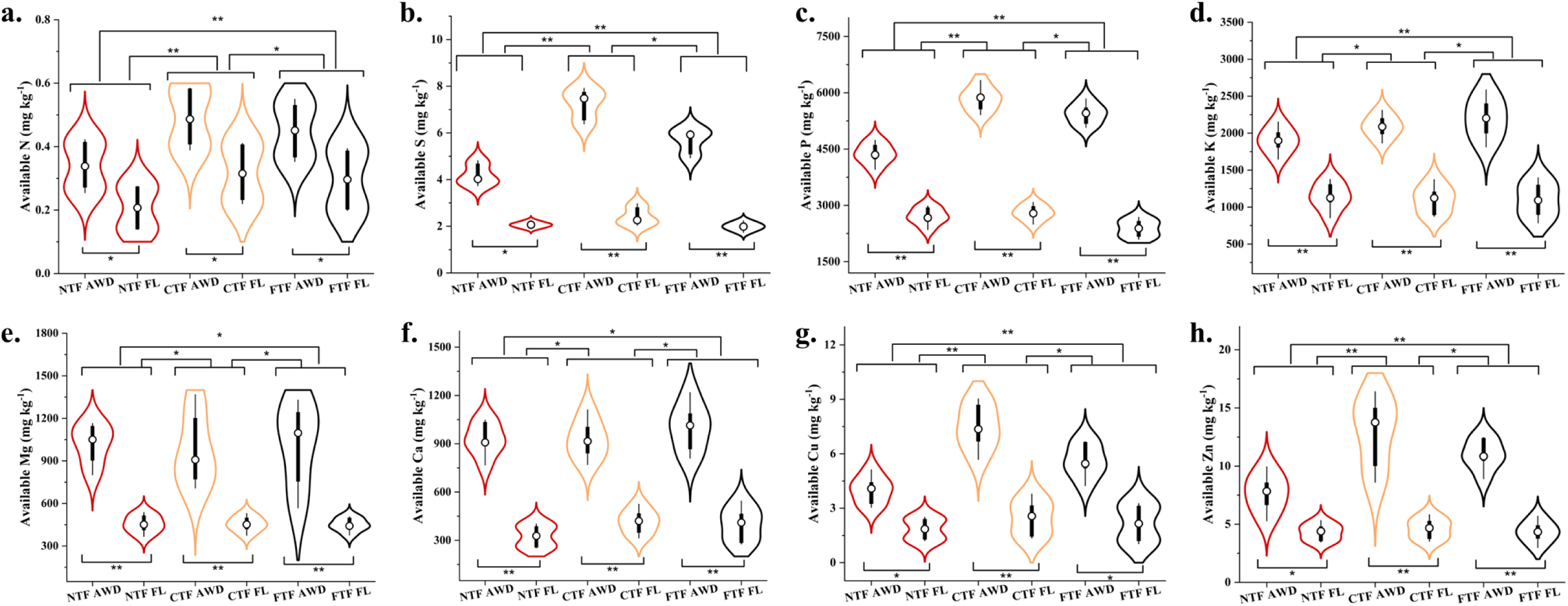
Bioavailable concentrations of selected soil elements N-S-P-K-Mg-Ca-Cu-Zn **(a-h)** from three field setups with varied tillage practices. Violin-box distribution plots are presented here. These are the average data of two-year soil sequential extraction analysis from 12 experimental sites. Distinct concentration differences and statistical significance variance have been found. ANOA with *p*<0.05 and *p*<0.01 significance levels are marked as ‘*’ and ‘**’, respectively, when compared within groups.

### Microbial diversity and metabolic responses

The microbial community is one of the primary bio-indicators of soil health and fertile lands. Tillage practices can drastically modulate soil microbial diversity and their molecular responses^29^. **Extended figure 14** shows the microbial relative abundance of phyla with the dominance of proteobacteria (30.3-39.4%), actinobacteria (20-43.1%), acidobacteria (8- 12.8%), chloroflexi (5-31.3%) and plactomycetes (2.9-7%). A significant reduction in microbial abundance was recorded while AWD was compared to FL practice under these three tillage practices. The fold change of relative abundance was reduced by 23.3%, 44.8% and 13.2%, respectively, in NTF, CTF, and FTF flooded irrigation compared to the AWD irrigation. The combined effect of tillage and irrigation was also evident in microbial metabolism, as shown in **Fig. 5**. Results from the high throughput sequencing showed the comparative and gradual increase in microbial soil metabolic activities in the order of CTF>FTF>NTF, as presented in **Fig. 5a**. This chord chart shows the read count-based separation of metabolic activities that are specifically expressed under the linked practice setups. Microbial metabolisms are pathway-dependent and understanding of these essential metabolic pathways through next-generation sequencing proved a higher occurrence and activity in CTF and FTF compared to NTF, as shown in **Fig. 5b**. This colored heat map shows the degree of occurrence of twenty essential metabolic pathways under AWD and Fl irrigation regimes were varied significantly, however, the tillage practice had distinct induction on these changes. The highest to lowest biological functioning of active pathways was traced in the order of CTF AWD > FTF AWD > FTF FL ≥ CTF FL > NTF AWD > NTF FL. Interestingly, under the developed FTF tillage with the AWD combination, environmental adaptation, metabolism of non-essential amino acids and other environmental biodegrading activities were triggered compared to the other tillage practices. This suggests the positive effect of this newly developed practice over the no-till system. These pathways were compared with the available database of the Kyoto Encyclopedia of Genes and Genomics (KEGG). Machine learning-based K-means cluster analysis showed that the microbial KEGG pathways can be analysed by lowering the dimensions using principal components and clustering. The generated data from the average points was further used here for a cross-check of the distribution pattern and reliability of the multiple factors through principal component analysis (PCA). The average data of KEGG reads were used to get PC1 with 99.89% variance and PC2 with 0.08% variance, as shown in **Fig. 5c**. Further clustering into NTF, CTF and FTF, the distance of each PC data point from the centroid showed the variability of microbial metabolism under these field setups. Shaded areas indicate the significance margin. From the figure, it is evident that the FTF-respective data are more variable with the highest distance values, whereas the lowest distance span is associated with the NTF and not within the range of the 95% confidence area. CTF data lies between these two sets. Panther pathways mostly indicate the active proteins that participate in two major domains- molecular functions and biological processes Metabolic activities that are linked with the biological processes and molecular functions are presented in Panther charts in **Fig. 5d**. From different categories listed in the figure as a pie graph, the catalytic activities under molecular functions while biological regulations and cellular processes under the biological processes were much lesser in NTF microbial populations than compared to the CTF and FTF population. Here, the Panther charts show the average data of AWD and FL to mark out the difference in results within tillage practices only. Interestingly, catalytic activity was found in much less in NTF compared to CTF and FTF, while biological regulation was higher in NTF than in CTF and FTF. The overall microbial metabolism under NTF, CTF and FTF under AWD and FL conditions was also tested using K-means cluster analysis, as shown in **extended figure 15**. The identified gene ontology (GO) terms from each of the tillage practices with irrigation regimes are presented in **extended figures 16** and **17** for high-frequency GO and low-frequency GO. Most of these high-frequency GO indicate the essential cellular synthetic and signalling pathway activities and respective biomolecules, whereas the comparatively low-frequency GO shows the structural features controlling activities and molecules. **Extended figure 18** provides the statistical insight into the correlative nature of microbial GO in all these field setups. This proves that although a difference was clearly found and established in this microbiota, a majority of the GO terms and metabolisms were present. This test was essential to prove that the degree of occurrence in KEGG or GO terms might differ, but the communal microbial metabolism remains active in all these fields.

**Fig. 5.**
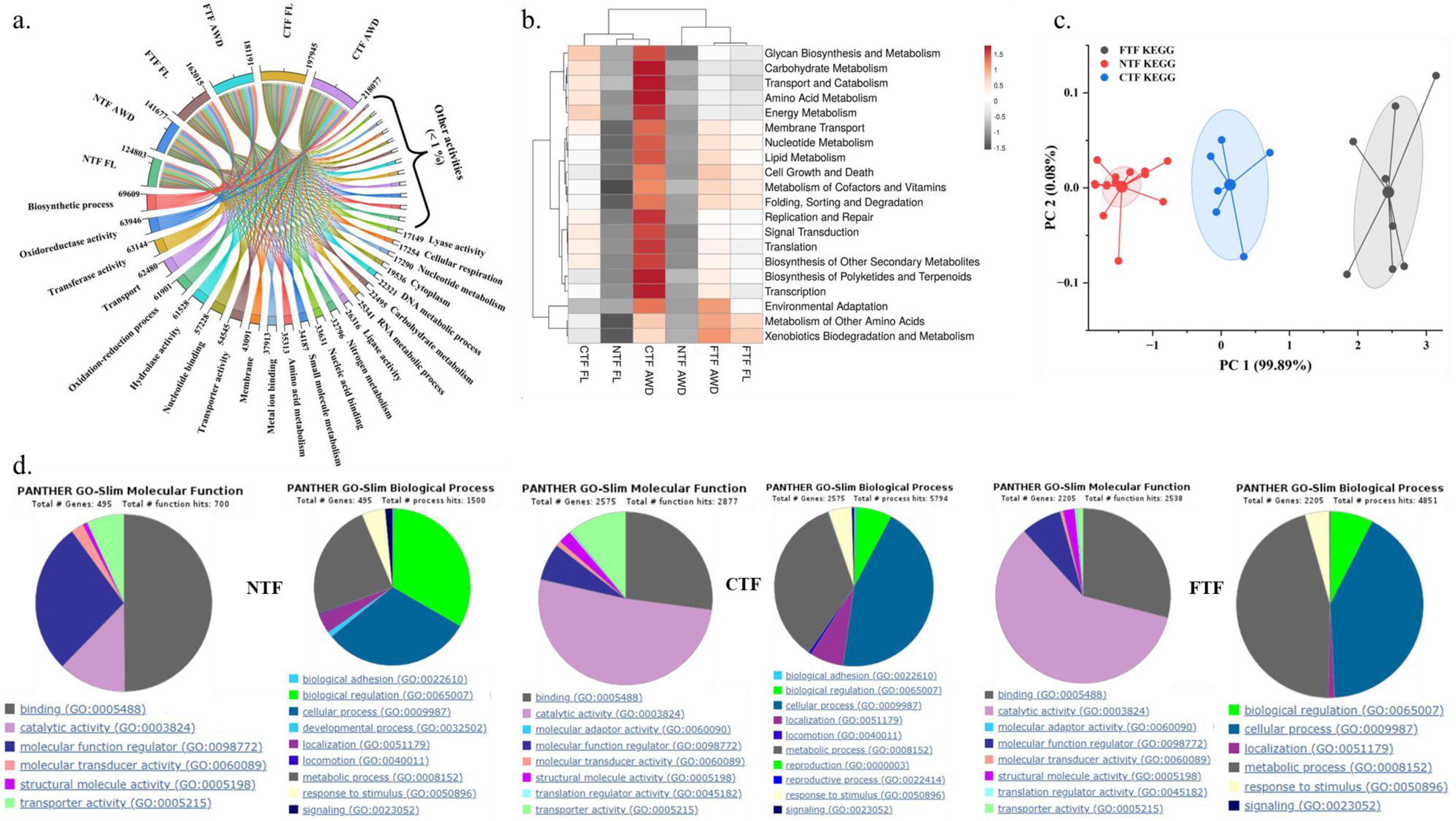
Microbial high-frequency metabolisms **(a)**, dominant KEGG pathways under differential setups **(b)**, comparative average KEGG reads using K-means cluster analysis with principal components **(c)**, and Panther pathway analysis of molecular functions and biological process-related GO-terms **(d)**. Data represents the average of 2 years of analysis for a triplicate data set.

### CO_2_ interactions and membrane transportation

When there is a CO_2_ flux in the soil, a variable rate of CO_2_ is released into the atmosphere, corresponding to the differential release of CO_2_ from microbial sources as part of their metabolism. This study uses experimental data and modelling tools to show the molecular interactions of CO_2_ molecules with the transporter proteins responsible for the passage through membranes. Carbon dioxide-binding proteins similar to the ammonium transporter (AmtB) and transporting hydrogenase proteins (HypC-HypD) are known for the passive passage of CO_2_ other than the regular diffusion process. **Fig. 6** shows the specific amino acid positions with the internal distances allowing comparatively smaller CO_2_ molecules to pass. In the AmtB protein, conserved histidine (His) residues at positions 168 and 318 are located in two separate α-strands with an internal gap of 4.42 Å, whereas the intra-atomic distance of CO_2_ is 2.4 Å, making it possible to pass through. Hydrogenase maturation factor HypC has a functional cysteine (Cys) at position 2 of its N-terminus, which closely interacts with a His at 51 position, maintaining a molecular distance of 7.81 Å and allowing CO_2_ to pass through. Another Fe- (CN)_2_CO cofactor assembly scaffold protein HypD forms the CO_2_ passage channel of 5.25 Å involving two Cys residues at positions 69 and 72 with a glutamic acid (Glu) at position 357. The models are presented here as a combination of Gaussian volume and Ribbon structure with targeted amino acids in ball-&-stick form to show the tunnel positions in these transporter proteins through which CO_2_ molecule passes. String networks of these Hya, Hyb, Hyp, Amt and Gln proteins show the inter-connections of related proteins that play roles in similar activities (**extended figure 19**). Depending on the species and route of CO_2_ passage, this network might get involved in all or some. Violet lines are experimentally proven interactions, whereas the cyan line and light green colors, respectively, indicate the database-curated linkage and literature-curated linkages of these protein molecules.

**Fig. 6.**
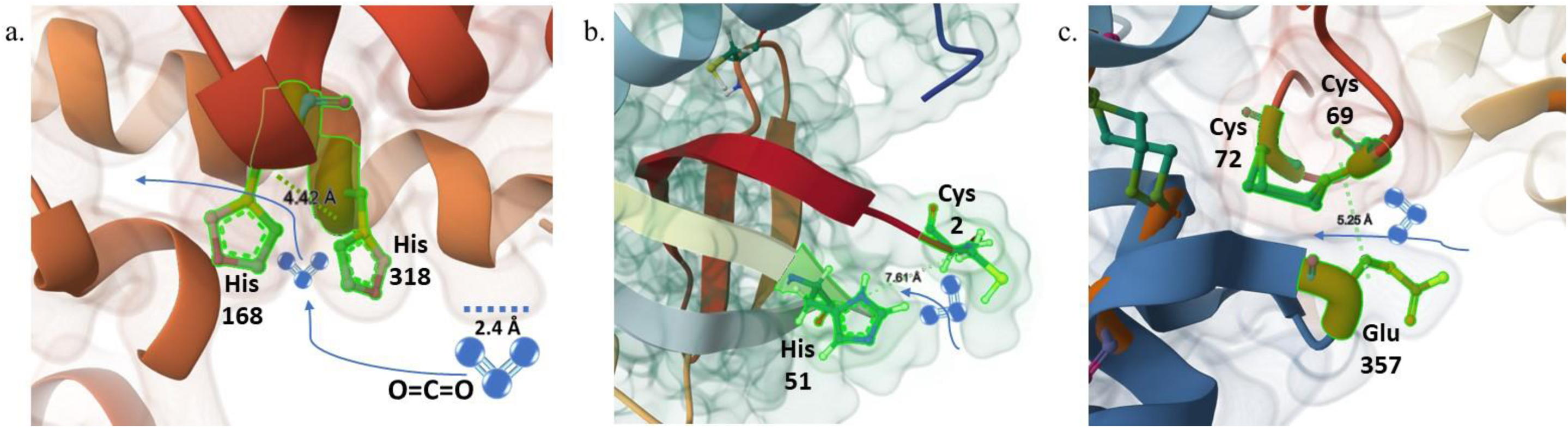
Localized carbon dioxide binding (**a.**, amtB) and transporting (**b.**, hypC, and **c.**, hypD) proteins with their modelled structures and bound CO_2_ within the core while mobilizing between inter-membrane spaces of the cell. CO_2_ is marked in yellow circle and thereafter in ball-&-stick structure. All the three proteins are shown in their structural ribbon forms with additional side groups in the ball-&-stick form.

## Discussion

### Efficiency of furrow tillage in CO_2_ release

Globally, the release of CO_2_ fluctuates depending on the associated sectors, although the annual and cumulative CO_2_ release map and the trend-line charts are indicative of a gradual increment in the CO_2_ content (**extended figures 20** and **21**). These maps and charts are based on the collective data and availability on the ‘Our World in Data’ site (https://ourworldindata.org/co2-and-greenhouse-gas-emissions). This change in land usage not only covers the anthropogenic invasion of the natural environment but also changes in the patterns of agricultural fields and practices. The cumulative data trend of CO_2_ release from 1993 to 2023 in all the continents shows the seriousness of this matter, directly linked to land-use change. The newly developed furrow tillage minimised the soil carbon release much less than the conventional deep tillage while maintaining the carbon pool similar to the no-tillage practice. This resulted from the controlled tilling of the side furrows and undisturbed soil beds. In a study by Sha et al^30^, the optimum land management in terrestrial vegetation has been analysed and shown to have a vital influence on the release of greenhouse gases. These previous studies prove the requirement of a proper agronomic practice that can balance soil carbon storage and loss while maintaining soil-crop qualities. In a study, it was experimentally shown and estimated that the SOC content and the plant carbon pool will not be equivalent when there’s an increase in the CO_2_ release in the atmosphere^31^. Also, there is an inverse relation between soil SOC and plant carbon pool based on plant nutrient accumulation preferences and microbial participation^32^. Some extensive global reviews are available that justify the point that following no-tillage has the least or no effect on maintaining or increasing the soil SOC stock^8,33,34^. This study experimentally provides evidence for the optimal status of SOC and CO_2_ released from the furrow tillage compared to the conventional deep tillage that releases CO_2_. While no-tillage has a conflict with the SOC stock maintenance compared to traditional tillage, it also reduces the soil aggregates and results in the soil micropores. Whereas reduced tillage or conventional tillage can have balanced micro to macroaggregates while soil pore sizes are also well connected^35^. The soil clods and water movement analysis in this study have proven the same by getting reduced vertical to lateral movement of water in NTF soils compared to the CTF and FTF. A report also suggests that the undisturbed soil bed in no-till soil can increase the soil diversity at first instance, but the gradual hardening of surface soil results in deprived O_2_ saturation in the soil, leading to reduced microbial metabolic activities^36^. Reports suggest that the tillage practice certainly improves nutrient mineralisation and availability; however, the release of CO_2_ increases depending on other agronomic practices, making it debatable^35,37^. This controversial state can be efficiently addressed by following the furrow tillage practice, which can minimise the soil carbon loss but maintain the nutrient availability to the plants. We adopted the OK method to predict the C release from places other than the experimental fields due to its suitability for interpolating and estimating the unobserved locations^38^. In the present study, the logarithmic variogram models were found to be the best-fit model in the OK method. Though the earlier work showed that the function of the best-fit variogram model can vary across the region^39,40^. Despite the fact that the OK method was carried out with the same number of data points in both years, the R^2^ values were found to be lower in the second year (2023). This might be attributed to the non-uniformity of the data values. The performance of kriging depends on the quality of the data^41^. In 2022, the data values were more uniform across the tillage practices as compared to those in 2023. However, the OK method was demonstrated as an accurate interpolation tool for carbon mapping due to high correlation with the field observations^39^. Further, Chabala et al^42^ found no significant difference between the observed and OK predicted values. Kılıç et al^43^ demonstrated that the OK method resulted in higher accuracy as compared to the inverse distance weighting (IDW), and empirical Bayesian Kriging (EBK) methods for estimating and mapping soil organic carbon. The accuracy of the OK method can be improved by integrating OK with multi-layered perception network method that reduces the underestimation of high values. The local carbon release maps generated from OK can be used a proxy for the carbon release from the fields other than the experimental plots. These maps can contribute to the optimisation of the tillage practices in the study region.

### Plant-microbial responses towards furrow tillage

Rice plants can efficiently express their stomatal movement under varied CO_2_ concentrations in the surroundings^44,45^. In this study, the time-dependent analysis of stomatal observation under scanning electron microscopy (SEM) shows visual proof of varied CO_2_ content in the surroundings of the sampled rice plants (Fig. 3). The released CO_2_ rate from NTF was the least whereas CTF had the highest release rate and the FTF remained as a median of these two, which was reflected in the open and closure state of the stomata. A plant’s nutrient uptake and transportation depend on the vasculature structures like the xylem and phloem that transport these molecules at distant locations, giving a proper nutrient distribution in plant tissues^46^. This, in turn, results in structurally confined vascular systems. Also, it has been shown that the elevated CO_2_ level can compromise xylem structure while prolonged opening of stomata due to low CO_2_ content can also hamper the guard cell activities^24^. This suggests the optimal CO_2_ release, as the FTF practice showed, can maintain the plant’s anatomical rigidity. Depending on the nutrient availability, CTF and FTF had a higher bioavailability compared to the NTF, resulting in better rigid xylem and phloem structures. Nutrient bioavailability is directly related to plant elemental uptake and total concentration^28^.

While no-till practice can increase the carbon stock and, hence, microbial biomass carbon at the preliminary phase, conventional tillage can increase nutrient mineralisation, create more labile carbon content for rapid microbial consumption, and increase metabolic activities. Furrow tillage, in this context, has optimum microbial cellular and metabolic activities. Results from high throughput sequencing show the active gene ontological terms and KEGG- associated pathways that are getting expressed more under the FTF practice compared to the no-till practice, giving us the choice of opting for the FTF method over the NTF. Although CTF showed the highest microbial activities, considering the fact of releasing least CO_2_ makes it less favourable. Soil microbial metabolism, gene expressions and protein activities vary with the conditional status of soil profiles. Reports are suggestive of the fact that deep tillage can influence functional gene expressions in microbial communities^47^, whereas the no-till practice can increase the carbon metabolising gene expressions^48^. In this study, the microbial communities from the furrow tillage have established proof of having a high cellular catalytic activity and structural molecule activity compared to the no-till microbial communities (as shown in the panther pathways). Essential metabolic pathway activities in FTF microbes, including biosynthetic processes, oxidoreductase activities, transference and hydrolase activities, metal ion binding, amino acids and nucleotide metabolism, cellular respiration, carbohydrate and lipid metabolism, cell growth regulator and environmental adaptation, are highly expressed similar to the CTF and greater than NTF. This indicates the active microbial gene responses under furrow tillage that can have benefits from both the NTF and CTF practices. CO_2_ released from the soil not only influences plant uptake through stomatal movement but also refers to microbial CO_2_ transportation and release. Studies based on experimental and simulator analytical assessments have shown the involvement of bacterial ammonium transporter-like AmtB protein in active CO_2_ passage^49,50^. Here, the structural features of amtB protein have been analysed using the updated AlphaFold and 3D viewer modelling software that shows the CO_2_ binding and interaction with His residues at positions 168 and 318 that create one of the active sites, as previously reported by Jung^51^. The other transporter proteins HypC and hypD are [NiFe]-hydrogenase accessory proteins that interact with and bind a Fe–(CN)_2_CO moiety and CO_2_ ^52^. HypC andHypD, separately and together as a complex, makes the passage for the CO_2_ molecule (Fig. 6). HypC delivers a Fe molecule, bound to a CO_2_ molecule, to HypD protein where the reduction reaction occurs, and a CO molecule generates. This suggests the process of continuous and active CO_2_ transportation in bacterial cells as a part of their carbon metabolism. These proteins work in a network of closely related proteins of HybG, HypE, HypF, alongside nitrogen metabolism controllers GlnB, GlnL and GlnG. The carbon dioxide transportation is a marker of bacterial cellular activity that can be strongly correlated with the overall microbial metabolism and relative abundance, which in turn depends on the status of the soil. This establishes the fact that following furrow tillage can provide the optimal benefit from both the no-till and conventional tillage practices in a long farmland practice.

So, to understand the exact status of CO2 retention and release from the agricultural fields, this newly developed furrow tillage practice can highlight benefits from both no-tillage and deep tillage practices. These thorough field trials elucidated limitations in both these established practices while indicating the efficiency of furrow tillage in optimal soil-C storage, greater nutrient availability that enhances plant physiology, triggers microbial metabolic activities and modulates carbon dynamics.

## Methods

### Field trials of the new tillage practice

According to the 2024-2025 report of the US Department of Agriculture (USDA), India is the highest rice-producing country in the world, contributing >27% of global production (https://www.fas.usda.gov/data/production/commodity/0422110). Hence, the Gangetic delta plain was selected for this study, conducted in the Nadia district of West Bengal, India. Variable tillage practices are often observed in India, dominating the conventional deep tillage. Therefore, sites were chosen where people follow both no-tillage and deep tillage practices. The fields within these experimental sites were randomly selected considering one constant: no-tillage field availability. So, the three field setups were easily compared during this 2-year-long experiment (2022 and 2023). The spatial coordinates of these 12 experimental sites are provided in the **extended table 1** and used in the geospatial mapping. Each of these fields was 800 m^2^ in area. The conventional tillage field (CTF) applies deep tilling using ploughing machines, while the no-tillage fields (NTF) are mostly undisturbed (Derpsch, 2003). We used a mechanical tilling tractor machine for an average of 20cm deep tillage. The proposed furrow tillage fields (FTF) were maintained regularly for optimum results. Each of the FTF had multiple furrows and beds, where a single bed had two-sided furrow channels. This FTF had two parts of field bed preparation- (i) the furrows were tilled using a hand plougher, keeping the depth 20cm as well, and (ii) the mid-bed remained undisturbed for the whole cultivation period. These furrows were tilled by this hand-plougher every 3 days. The schematic and field representations are shown in the **extended figure 1**.

### Soil clods and water movements

The soil surface was analysed using porosity analysis and field water movement tracking to understand any surface structure changes occurring during the tillage practices and irrigation regimes. The surface horizons from H1 to H3 were found to be in two categories- Г and Δ clods^53^. The top 0-4cm layer was highly porous in formation and included in the Δ clod, while the below soil structures to 15cm were mostly small aggregates and can be considered as Г clod. Below 15cm, the macro-aggregates were noticed. Soil clods are responsible for developing the soil pores that allows water passage through these pores. Fields with NTF, CTF and FTF setups were sampled for both vertical and horizontal cores to check the soil porosity. Vertical coring was restricted in 0-20cm whereas horizontal cores were collected within 0-5cm. collected cores were incubated under a vacuum condition to minimize any mechanical structural damage. The bulk density and particle density were then measured following the method by Sasal et al^54^. These two porosities were used to define the total porosity in soil. Another measurement was carried out following this method, termed as structural porosity. Finally soil textural porosity was measured as the difference between total porosity and structural porosity. This textural porosity helps to understand the water holding and movement status under differential tillage practices. Thus, measurements of soil clods helped us define the flow system linked to the soil structures in the mid-bed and the tilled furrows as shown in **extended figure 2**.

### Irrigation regimes and rice cultivation

Irrigation regimes were essential to consider representing the exact data from the field trials, as they play a crucial role in soil elemental dynamics and GHG release. With the three field setups of NTF, CTF, and FTF, the study also implemented conventional flooding methods and deficit irrigation. Alternate wetting-drying (AWD) and flooded lands (FL) were maintained for each of the three tillage forms. So, six field setups were established in all 12 experimental sites. For flooding, all the fields were submerged under a 10cm waterlog for the first month and a 30-35cm waterlog was maintained till the late-inflorescence stage of rice cultivation. However, it is essential to mention that the AWD irrigation in the FTF slightly differed from the NTF and CTF. The AWD practice was followed by the method developed by Majumdar et al^55^ for both CTF and NTF, which kept the waterlog condition for 2 days, followed by the drying phase of 4 days. In FTF, this pattern was modified by retaining the waterlog for 2 days in the whole field, followed by a drying phase for the mid-bed only. Furrows were always filled with water during the drying phase.

### Measurements of labile carbon and released carbon dioxide

Labile soil carbon contains three main types: potassium permanganate (KMO_4_) oxidizable carbon (KOC), water extractable carbon (WEC), and microbial biomass carbon (MBC). Soil samples were mixed up with 40mM KMO_4_ and shaken for 6-7 hrs followed by a centrifugation. The collected supernatant was then measured at the OD of 565 nm. Another batch of soil samples was water-extracted following a shaker and centrifuge, which thereafter mixed up with 0.07 N potassium dichromate (K2Cr2O7), sulphuric acid (H2SO4) and ortho-phosphoric acid (H3PO4). This mixture was heated at 120°C for 2 hrs, and the extracted supernatant was titrated using a 0.035 N ferrous ammonium sulphate solution^56^.

For the released CO_2_ from fields, specially designed fibre boxes were used to entrap the gaseous release from the soil, as shown in the **extended figure 3**. The released CO_2_ was collected using a syringe attached to a gas collection bag with a stopper. Collected samples were then taken to the lab for a gas chromatography mass spectrometry (GC-MS, Agilent 8890) analysis. The instrumental specifications are followed as per the setup described by Ussiri and Lal^57^. This facility was equipped with a 3m long and 0.3m diameter width Hayesep D column and a thermal conductivity detector. The temperatures of the oven and the detector were 55°C and 160°C, respectively. Inert helium was used at the rate of 20cm^3^ min^-1^ as a carrier gas. For the internal calibration, CO_2_ standard gas was used. The below equation was used for the assessment of daily CO_2_ flux rate-

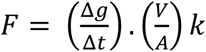

Where F is the flux rate (g CO_2_-C m^2^ d^-1^), CO_2_ concentration change in the linear form inside the chamber (Δg/Δt), *V* and *A* are the chamber’s volume and surface area, respectively, and *k* is the conversion factor of the analysis time. Time vs concentration data of CO_2_ flux rate per hour was calculated using the linear regression model.

### Spatial distribution mapping of released CO2 from the field

Based on the experimental data local CO_2_ release map was generated using geostatistical interpolation technique. Ordinary Kriging (OK) method was adopted to estimate the values of the CO2 released from the areas other than the experimental fields. The mean of a stationary variable is assumed to be a constant in the local neighborhood, and the semi-variance changes with the distance only not with the location. The best-fitting semi-variogram function is used in kriging to predict the spatial variability of a random variable. A linear predictor is adopted in OK method with the following formula (https://bookdown.org/yan_ren0518/summary_note/spatial-prediction-and-kriging.html)

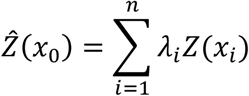

Where, *Ẑ*(*x*_0_) is the estimated value at location *x*_0_; *Z*(*x*_𝑖_) is the observed value at sampled location *x*_𝑖_; 𝜆_𝑖_ is the weight assigned to each observed value and n is the number of sampled points used for estimation. The sum of weights i.e. 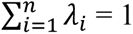.

In the present study, the “Ordinary Kriging” tool under the “Geoprocessing” of SAGA (System for Automated Geoscientific Analysis) 6.4.0 was employed for OK of the CO_2_ released from the experimental fields. Earlier, scientists demonstrated the efficiency of OK method for spatial interpolation^22,23^. The best-fit variogram models obtained in the present study have been presented in the **extended table 1**. In order to represent global insight on the carbon emission during the experimental year, the Global Gridded Daily CO2 Emissions Dataset (GRACED) data were used in the present study. GRACED provides near-real-time global gridded daily CO_2_ emission data at 0.1° X 0.1° spatial resolution in kg C per hour unit^58^. Spatial emission data from several sources, including near-real-time daily national CO_2_ emissions estimates (Carbon Monitor), were used to develop the GRACED dataset. The GRACED dataset has already been proven as a reliable data source with acceptable uncertainty^59^. We calculated monthly carbon emissions from the daily datasets using the “Raster Calculator” of Quantum Geographic Information System (QGIS) 3.16.8. Global maps representing monthly average C emission (kg C/ hour) for 2022 and 2023 were generated in QGIS 3.16.8. This will differentiate the degree of C-release from other sources compared to the agricultural practices.

### Microbial biomass and soil physico-chemical properties

Soil physico-chemical properties were analysed to determine the correlative features of pH, redox potentials and microbial biomass carbon with SOC and LC. Oakton Waterproof PCS Testr 35 and ORP Testr 10 were used for measuring in-field soil pH and redox potentials, respectively. The data were collected from 20 spots from each field setup, followed by developing a contour map of soil SOC and LC distribution patterns. Similarly, soil samples from the same spots were collected in Eppendorf tubes and stored in ice for microbial biomass carbon analysis. The traditional chloroform fumigation extraction method was followed with minor modifications (Witt et al., 2000). Briefly, the moist soil samples were kept under a vacuum for a 5-day incubation at 37°C with concentrated chloroform. After the incubation period, all the samples were extracted with potassium permanganate, followed by titration using acidified ferrous ammonium sulfate. The data of biomass carbon from fumigated to non-fumigated samples were tallied.

### Total and bioavailable elemental analysis with distribution patterns

Soil and rice plants were treated with different chemicals for the total and bioavailable fraction analysis of major soil elements that have some effects on the plant growth. Here, the elements considered are nitrogen, sulfur, phosphorus, potassium, magnesium, calcium, copper, and zinc. Total nitrogen and sulfur were determined by the Kjeldahl and turbidimetric methods, respectively. Whereas other elements were analysed for the total concentration using wavelength dispersive X-ray fluorescence (WD-XRF, S8 Tiger, Bruker) spectrophotometer^60^. The NIST 2711a, SDC-1 and JSD-2 were used as standard reference materials (SRM) for the internal calibration of the XRF. NIST 2711a was also used to calibrate an inductively coupled mass spectrophotometer (ICP-MS, Perkin Elmer) during the bioavailable elemental distribution. Soil samples were treated following a five-round extraction process that is called the sequential extraction method^61^. The first two fractions correspond to the water-soluble and ionic change concentrations and are jointly considered to be the bioavailable form. Plant samples were analysed for the total elemental concentrations following a di-acid digestion-dilution method^20^ . Rice flour NIST 1568b was used as the SRM for plant sample calibration.

### Plant physiology and anatomical observations

Plants growing on the fields were tested for the stomatal structure on the leaf surface and also for the internal ultra-structure of the vascular system (xylem, and phloem). Stomatal opening and closing depend on the availability of CO2 in the surrounding air. A leaf was marked at the start of the experiment (0th hr), and after 2 hrs of stand time, the same leaf was sampled again. A clean scissor was used for the 1^st^ cut from the top leaf part and placed it in the liquid nitrogen for a snap-freezing followed by a storage in dry-ice box while transporting to the lab. After 2 hrs, the same leaf was again cut from the middle part and snap-frozen. The schematic representation can be found in the **extended figure 4**. For the ultra-structure of the leaf vascular system, another leaf of the same plant was considered. An ultra-microtome was used for thin sectioning. Later, these cuts were visualized under the field emission scanning electron microscope (FE-SEM, Zeiss Supra 55VP) following the method by Majumdar et al^62^.

### Microbial community structure and next-generation sequencing

Fresh soil samples from the fields were collected in a dry-ice box and stored for further analysis. To isolate the DNA from these soil samples, a Qiagen DNeasy Power soil kit (Cat# 1288-50) was used and processed for the high throughput sequencing. The extracted DNAs were further eluted in nuclease-free water (Qiagen Cat# 129115), followed by a quality check using a nanodrop (Thermo Scientific Nanodrop Lite). Further, with all the qualified DNA samples, a library was prepared using ligation sequencing and PCR barcoding kits (Oxford Nanopore Technologies, SQK-LSK109 and EXP-PCR096). Amplicon DNAs were further end-repaired using the NEBnext Ultra II end repair kit (New England Biolabs, MA), while AmPure bead (Beckmann Coulter, USA) was used for further cleaning of the DNA clutters. The barcoded DNA was pooled using a dA-Tailing Module mixed with the NEBnext ultra II end repair kit. To get the optimal purity, another cleaning set was done following the NEB blunt/TA Ligase kit manual (New England Biolabs, MA). Oxford Nanopore Technology was used for the final round of sequencing on GridION X5 with a flow cell R9.4 SpotON version (FLO-MIN106). Nanopore fast5 format was used for the raw reads to make the base-called and further demultiplexed using Guppyv2.3.4. The OTUs and other processed reads were then checked for the community diversity indices, gene ontological terms, and metabolic pathway components through Kyoto Encyclopaedia of Genes and Genomes (KEGG), CO2-responsive gene networking and CO2 transporting gene product analysis.

### Molecular data set and CO2 transporter modelling

The curated data sets from the high throughput sequencing were divided into three categories-gene ontological terms, panther pathway terms and KEGG terms. These terms were marked according to their read counts from the NGS records and segregated depending on the NTF, CTF and FTF classes. A cord diagram was generated for the gene ontological terms showing the interconnections of the major metabolic pathway markers linked to these three field setups with respective read counts. A heatmap was generated following the chord plot to show the occurrence patterns and detected levels of major and minor gene ontology terms in these three field-microbial communities. Panther pathways are more related to the proteins that participate in diverse metabolic and cellular activities and shown here to identify what are the primary cellular activities getting changed within these microbial communities. KEGG metabolic pathways and their related terms were also presented as a heatmap to show the degree of occurrence of these metabolic markers or major pathway directories that got diverse values from this NGS sequencing.

Microbial CO_2_ transporter proteins are not exclusive to this gaseous molecule and primarily transported by hypD+hybG complex and amtB protein. The genes for these proteins were detected in most of the aerobic bacterial communities from the three fields; however, their degree of expression was different, and that led to research on the molecular structure prediction of these proteins and interactions with a passing CO_2_ molecule. AlphaFold 3, NCBI database and RCSB PDB 3D Mol Viewer were used to generate the predictive model of these membrane transporters with CO_2_-bound conditions. The protein sequence for each of these transporters was taken from the NCBI and loaded in the AlphaFold 3 to generate the model with a high Predicted Local Distance Difference Test (pLDDT) ≥85 scores. This selected model version was then further analysed in the 3D Mol Viewer to understand the locations of active sites where CO_2_ binds during the trans-membrane passage. For a better visualization and understanding, bulk protein chains were shown in Cartoon and Gaussian volume whereas the targeted amino acids were shown in ball-&-stick with element index coloring for the whole model. Internal distances between two targeted amino acids were determined using the Mol Viewer’s measurement tool and compared with the diameter of a CO_2_ molecule.

### Statistical analyses and data presentation

Statistical analyses were an integral part of this study, checking variability and significance level of each of the data sets. All the soil carbon content, released CO_2_, total and bioavailable soil-plant elemental concentrations were tested for the statistical significance at *p*<0.05 and *p*<0.01 levels using the analysis of variance (ANOVA). Further statistical justifications were checked by using four post-hoc analyses-Bonferroni correction, Mann-Whitney U test, Tukey honestly significant difference (HSD) test and Fisher’s least significant difference (LSD) test. All the details of these tests are shown in the extended figure 7 and extended tables 2-4. Linear regression analysis was also tested for soil carbon data and microbial GO distribution. The molecular responses from microbial communities recorded from different soil setups were tested with the machine learning-based K-means cluster analysis to identify the principal components and variable range within the 95% significance area. All the data sets used for these analyses in this study were representative of the 2-year average data. Origin 2024, Graphpad Prism var. 6, SigmaPlot var. 12, and Shiny GO online var . 0.77 were used.

## Acknowledgement

This work was funded by the National Postdoctoral Fellowship scheme, Ministry of Education, Government of India, with a project number PDF/2022/001418/LS and Marie Skłodowska- Curie-UKRI Postdoctoral Fellowship scheme, United Kingdom, with a File number 101152605 and Council reference-EP/Z002664/1. A.M. acknowledge the support from Imperial College London Library during this study and funding for the publication.

## Authors contribution

**A.M.** conceptualised the project, received funding, performed experiments, and wrote the manuscript; **M.K.U.** managed fields and performed partial experiments with analytic measurements; **A.G.** did the data curation and spatial distribution mapping; **R.B.** did the CO2 measurements and analysis; **I.K-L., M.B.** and **T.C.** supervised the projects and revised the manuscript; **M.T.** helped in project development, ideological inputs and manuscript write-up; **D.M.** helped in statistical analysis and partial write-up; **B.G.** and **M.K.J.** helped in instrumental setup and analysis, including XRF and ICP-MS.

## Ethical declaration and competing interests

Authors declare no competing interests and ethical violations.

