## Supplementary files for "Furrow tillage reduces soil carbon loss and enhances microbial metabolism"

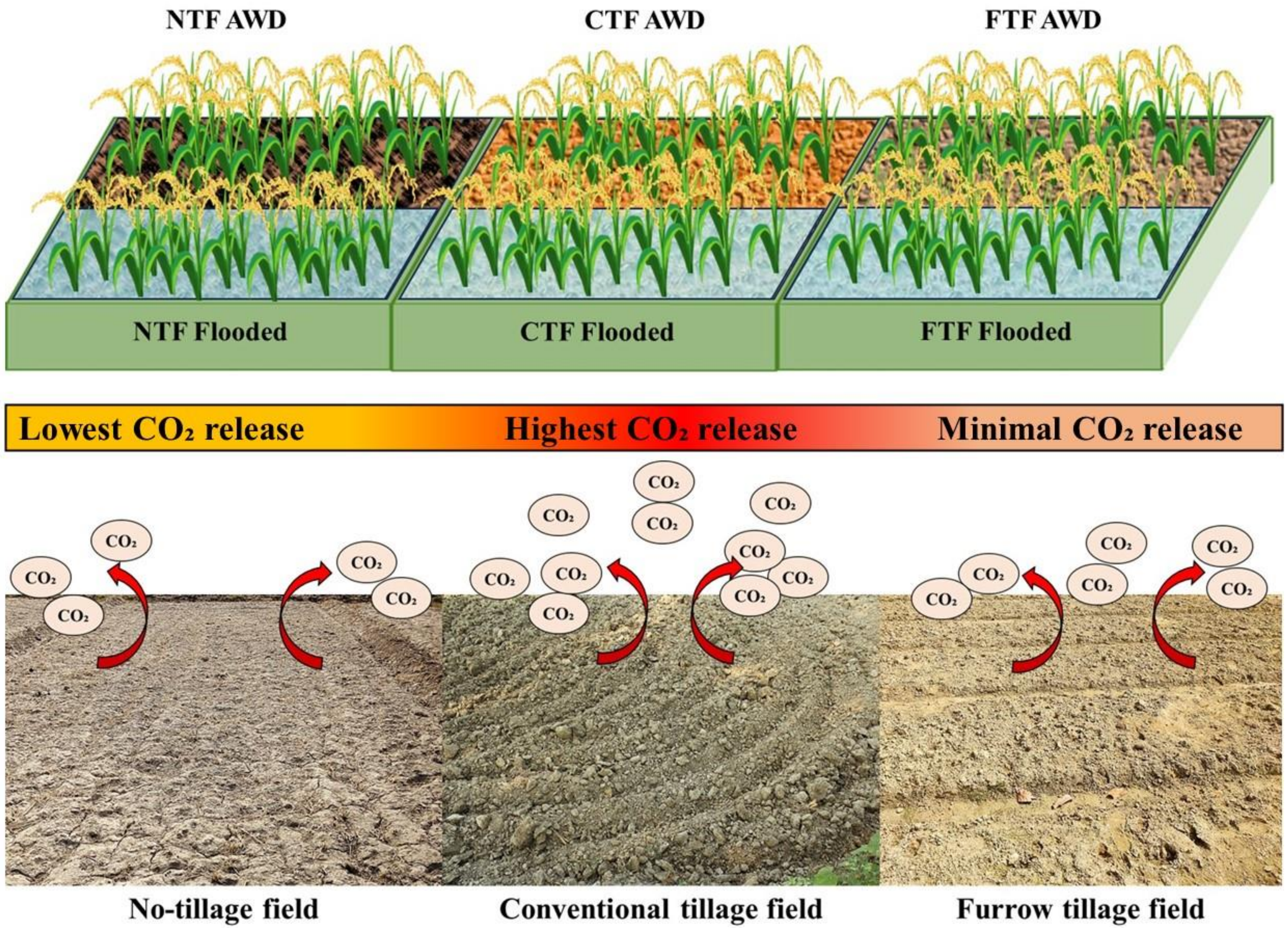

**Extended figure 1.** The three field setups are presented in a combination of schematic diagrams of irrigation regimes and varied tillage practices in the field. Released carbon dioxide ($CO_2$) bar is a colored illustration of analysed data during sampling phases within the two-year span.

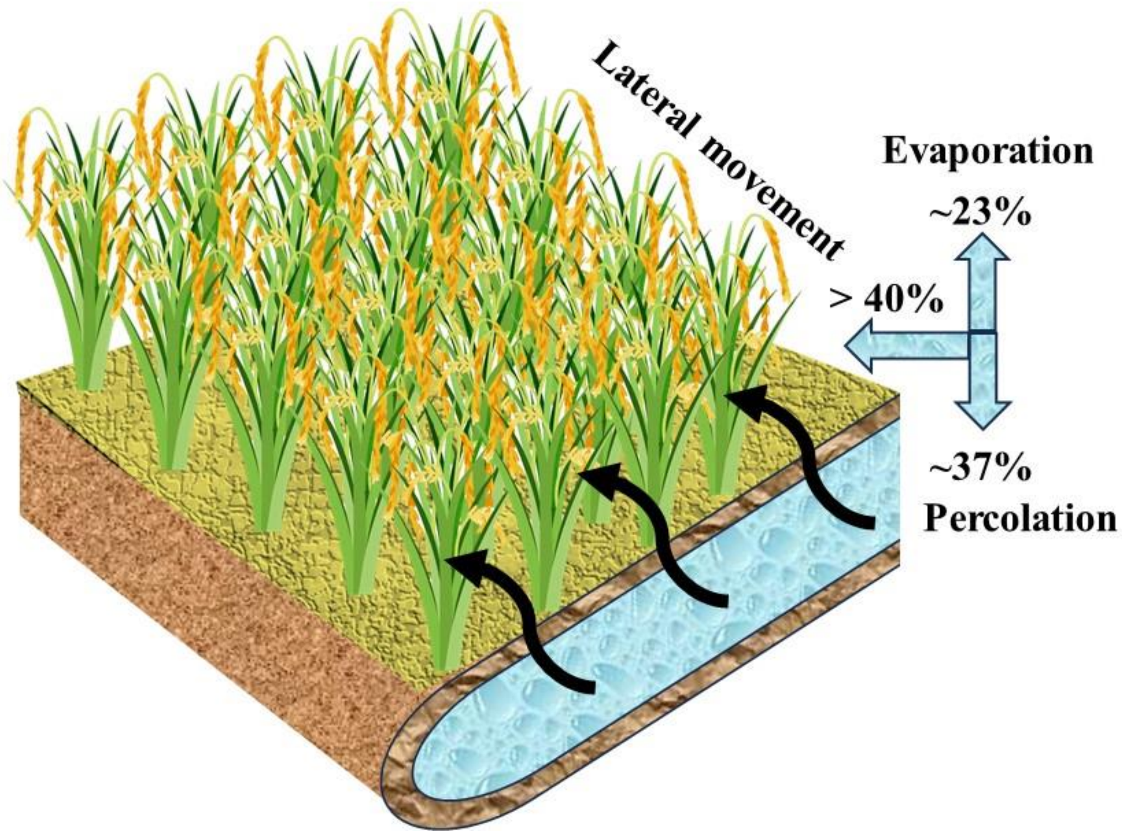

**Extended figure 2.** Furrow tillage water movement in three directions, as per the soil clods and water passage modelling.

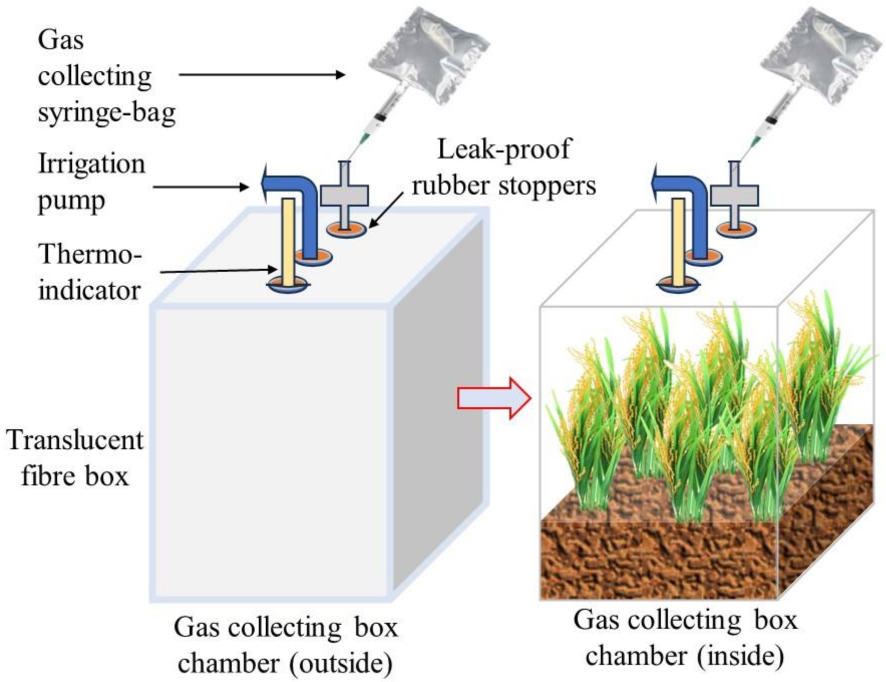

**Extended figure 3.** Schematic of a carbon dioxide gas collection chamber setup in the field.

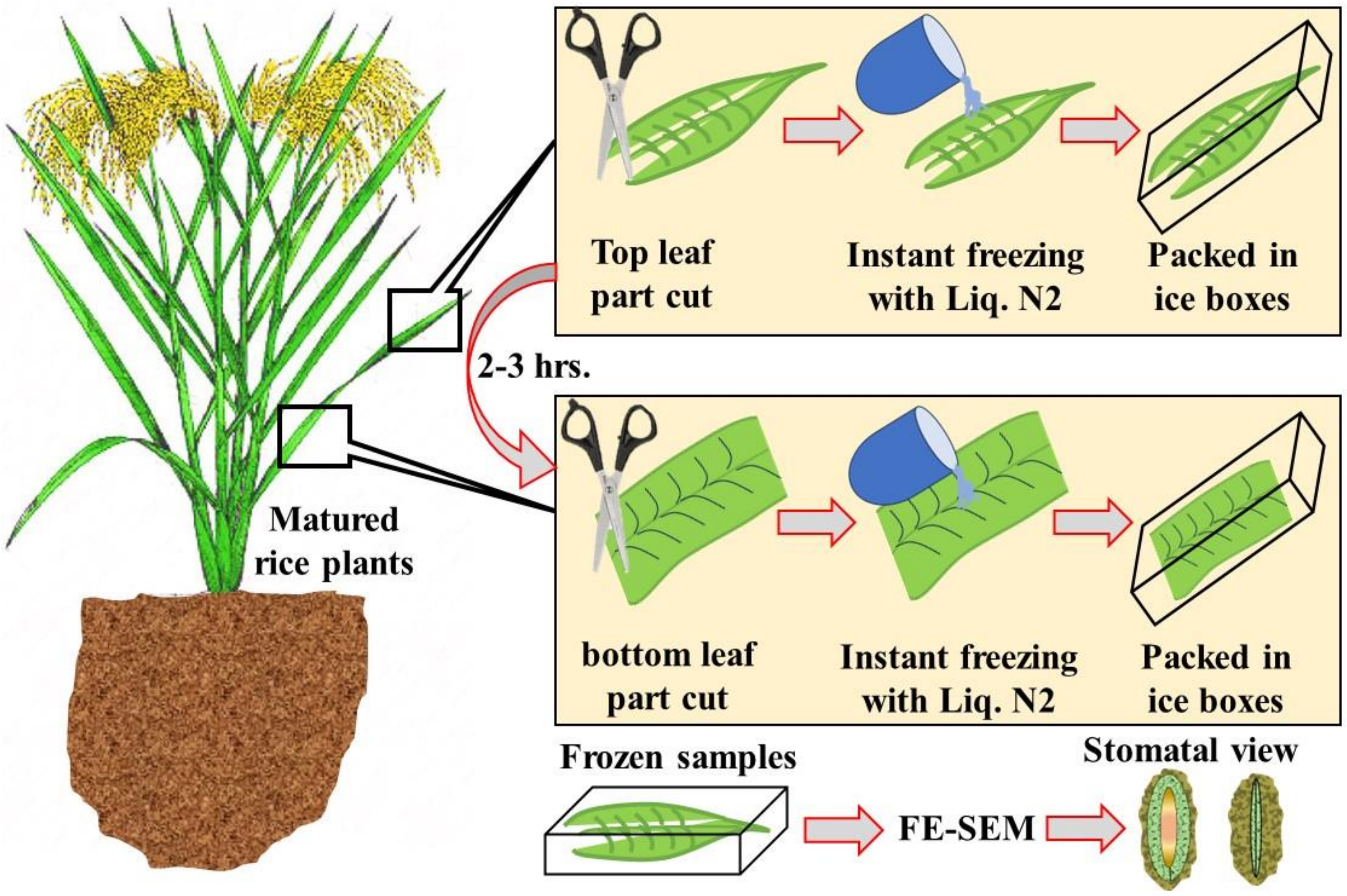

**Extended figure 4.** Plant leaf sampling and snap-freezing for internal structural observation.

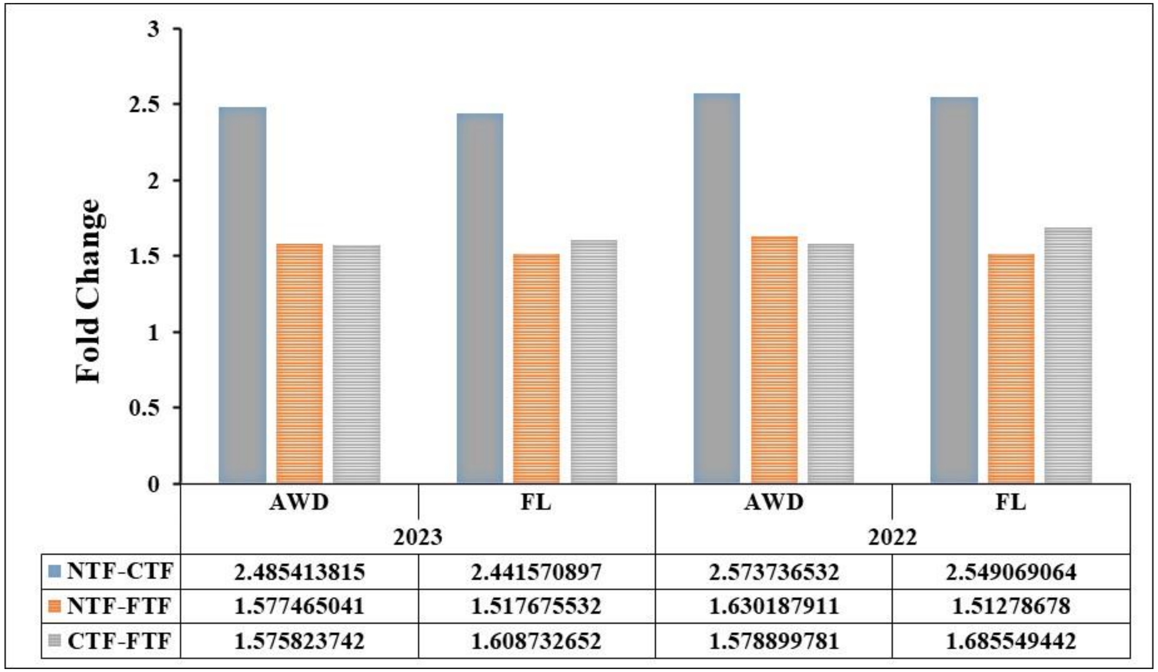

| | 2023 AWD | 2023 FL | 2022 AWD | 2022 FL |
|---|---|---|---|---|
| NTF-CTF | 2.485413815 | 2.441570897 | 2.573736532 | 2.549069064 |
| NTF-FTF | 1.577465041 | 1.517675532 | 1.630187911 | 1.51278678 |
| CTF-FTF | 1.575823742 | 1.608732652 | 1.578899781 | 1.685549442 |

**Extended figure 5.** Fold change of released $CO_2$ from three field setups in two consecutive years. The data represents the average trend results from the 12 experimental sites. This data has been further justified by one-way ANOVA at $p<0.05$ significance level.

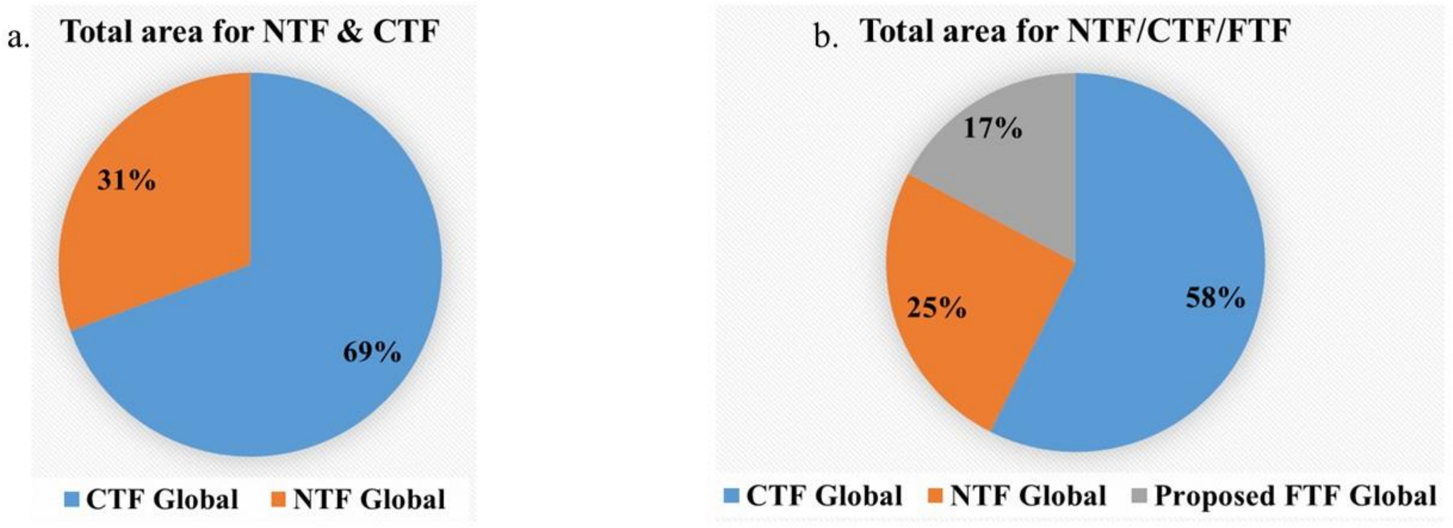

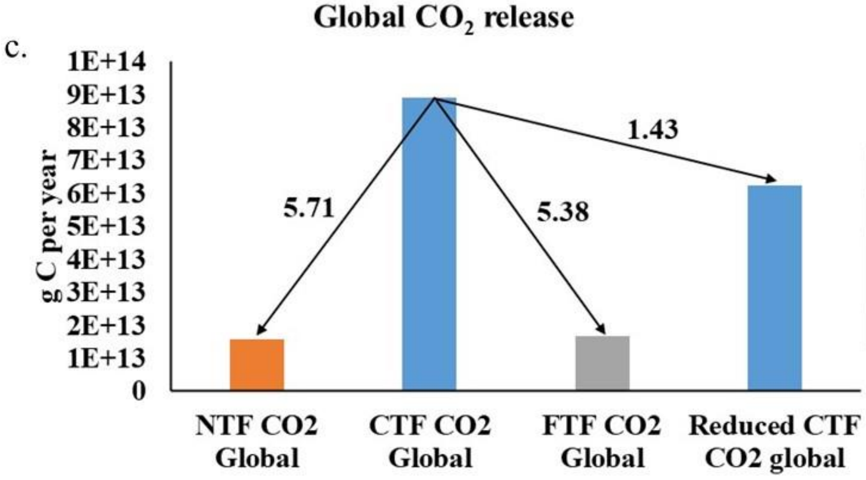

| Global CTF (hec.) | Global NTF (hec.) |
|---|---|
| 1150000000 | 507000000 |
| Reduced global CTF (hec.) | Proposed global FTF (hec.) |
| 805000000 | 345000000 |

**Extended figure 6**. Global agronomic field area for NTF and CTF with proposed FTF area coverage(a-b), and fold change in $CO_2$ release if FTF is followed globally (c).

a. Stats of SOC

*Means Comparisons*

*Bonferroni Test*

| | MeanDiff | SEM | t Value | Prob | Alpha | Sig | LCL | UCL |
|---|---|---|---|---|---|---|---|---|
| CTF NTF | -7.0726 | 0.66552 | -10.62723 | <0.0001 | 0.05 | 1 | -8.75118 | -5.39402 |
| FTF NTF | -3.5626 | 0.66552 | -5.35313 | <0.0001 | 0.05 | 1 | -5.24118 | -1.88402 |
| FTF CTF | 3.51 | 0.66552 | 5.2741 | <0.0001 | 0.05 | 1 | 1.83142 | 5.18858 |

*Tukey Test*

| | MeanDiff | SEM | q Value | Prob | Alpha | Sig | LCL | UCL |
|---|---|---|---|---|---|---|---|---|
| CTF NTF | -7.0726 | 0.66552 | 15.02917 | <0.0001 | 0.05 | 1 | -8.70562 | -5.43958 |
| FTF NTF | -3.5626 | 0.66552 | 7.57047 | <0.0001 | 0.05 | 1 | -5.19562 | -1.92958 |
| FTF CTF | 3.51 | 0.66552 | 7.4587 | <0.0001 | 0.05 | 1 | 1.87698 | 5.14302 |

*Fisher Test*

| | MeanDiff | SEM | t Value | Prob | Alpha | Sig | LCL | UCL |
|---|---|---|---|---|---|---|---|---|
| CTF NTF | -7.0726 | 0.66552 | -10.62723 | <0.0001 | 0.05 | 1 | -8.4266 | -5.7186 |
| FTF NTF | -3.5626 | 0.66552 | -5.35313 | <0.0001 | 0.05 | 1 | -4.9166 | -2.2086 |
| FTF CTF | 3.51 | 0.66552 | 5.2741 | <0.0001 | 0.05 | 1 | 2.156 | 4.864 |

b. Stats of labile C

*Means Comparisons*

*Bonferroni Test*

| | MeanDiff | SEM | t Value | Prob | Alpha | Sig | LCL | UCL |
|---|---|---|---|---|---|---|---|---|
| CTF NTF | 0.15299 | 0.00814 | 18.78726 | <0.0001 | 0.05 | 1 | 0.13245 | 0.17353 |
| FTF NTF | 0.10168 | 0.00814 | 12.48596 | <0.0001 | 0.05 | 1 | 0.08114 | 0.12221 |
| FTF CTF | -0.05131 | 0.00814 | -6.30129 | <0.0001 | 0.05 | 1 | -0.07185 | -0.03077 |

*Tukey Test*

| | MeanDiff | SEM | q Value | Prob | Alpha | Sig | LCL | UCL |
|---|---|---|---|---|---|---|---|---|
| CTF NTF | 0.15299 | 0.00814 | 26.56919 | <0.0001 | 0.05 | 1 | 0.13301 | 0.17297 |
| FTF NTF | 0.10168 | 0.00814 | 17.65782 | <0.0001 | 0.05 | 1 | 0.08169 | 0.12166 |
| FTF CTF | -0.05131 | 0.00814 | 8.91137 | <0.0001 | 0.05 | 1 | -0.07129 | -0.03133 |

*Fisher Test*

| | MeanDiff | SEM | t Value | Prob | Alpha | Sig | LCL | UCL |
|---|---|---|---|---|---|---|---|---|
| CTF NTF | 0.15299 | 0.00814 | 18.78726 | <0.0001 | 0.05 | 1 | 0.13642 | 0.16956 |
| FTF NTF | 0.10168 | 0.00814 | 12.48596 | <0.0001 | 0.05 | 1 | 0.08511 | 0.11824 |
| FTF CTF | -0.05131 | 0.00814 | -6.30129 | <0.0001 | 0.05 | 1 | -0.06788 | -0.03475 |

c. Stats of released CO2

*Means Comparisons*

*Bonferroni Test*

| | MeanDiff | SEM | t Value | Prob | Alpha | Sig | LCL | UCL |
|---|---|---|---|---|---|---|---|---|
| CTF NTF | 4.00667 | 0.0783 | 51.16824 | <0.0001 | 0.05 | 1 | 3.80917 | 4.20417 |
| FTF NTF | 1.47833 | 0.0783 | 18.87946 | <0.0001 | 0.05 | 1 | 1.28083 | 1.67583 |
| FTF CTF | -2.52833 | 0.0783 | -32.28878 | <0.0001 | 0.05 | 1 | -2.72583 | -2.33083 |

*Tukey Test*

| | MeanDiff | SEM | q Value | Prob | Alpha | Sig | LCL | UCL |
|---|---|---|---|---|---|---|---|---|
| CTF NTF | 4.00667 | 0.0783 | 72.36282 | <0.0001 | 0.05 | 1 | 3.81453 | 4.19881 |
| FTF NTF | 1.47833 | 0.0783 | 26.69959 | <0.0001 | 0.05 | 1 | 1.28619 | 1.67047 |
| FTF CTF | -2.52833 | 0.0783 | 45.66323 | <0.0001 | 0.05 | 1 | -2.72047 | -2.33619 |

*Fisher Test*

| | MeanDiff | SEM | t Value | Prob | Alpha | Sig | LCL | UCL |
|---|---|---|---|---|---|---|---|---|
| CTF NTF | 4.00667 | 0.0783 | 51.16824 | <0.0001 | 0.05 | 1 | 3.84736 | 4.16598 |
| FTF NTF | 1.47833 | 0.0783 | 18.87946 | <0.0001 | 0.05 | 1 | 1.31902 | 1.63764 |
| FTF CTF | -2.52833 | 0.0783 | -32.28878 | <0.0001 | 0.05 | 1 | -2.68764 | -2.36902 |

**Extended figure 7**. Post-hoc statistical analyses after one-way ANOVA of SOC, labile carbon content and released $CO_2$ from fields using Bonferroni correction, Tukey HSD, and Fisher LSD tests. The red-lined box marked scores of 1, indicating statistical significance.

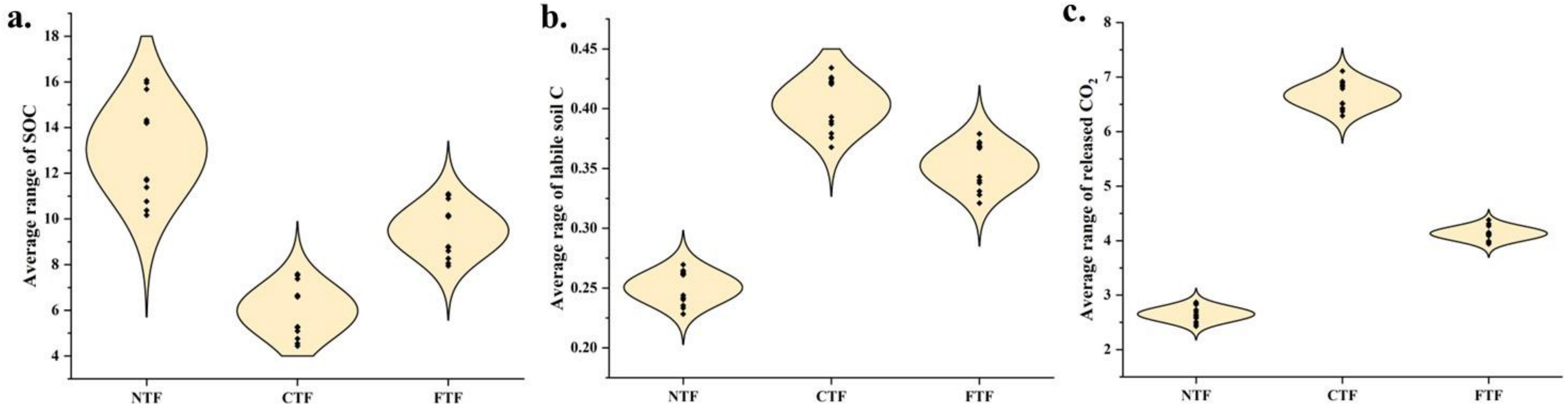

**Extended figure 8**. Average data trend of SOC, labile soil carbon and released $CO_2$ in the normal distribution model.

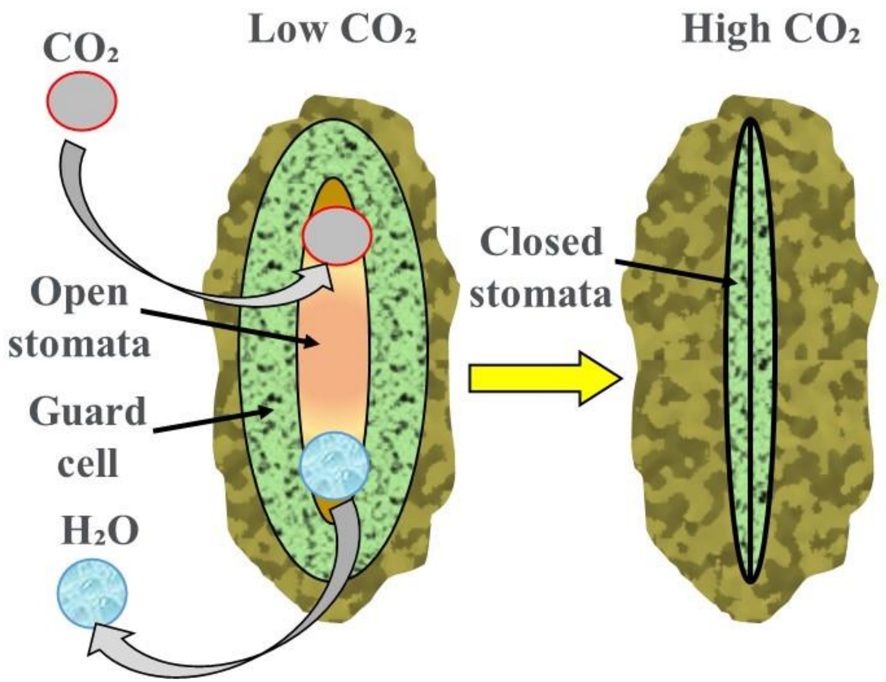

**Extended figure 9.** Schematic of a plant leaf stomatal activity under low and high $CO_2$ availability.

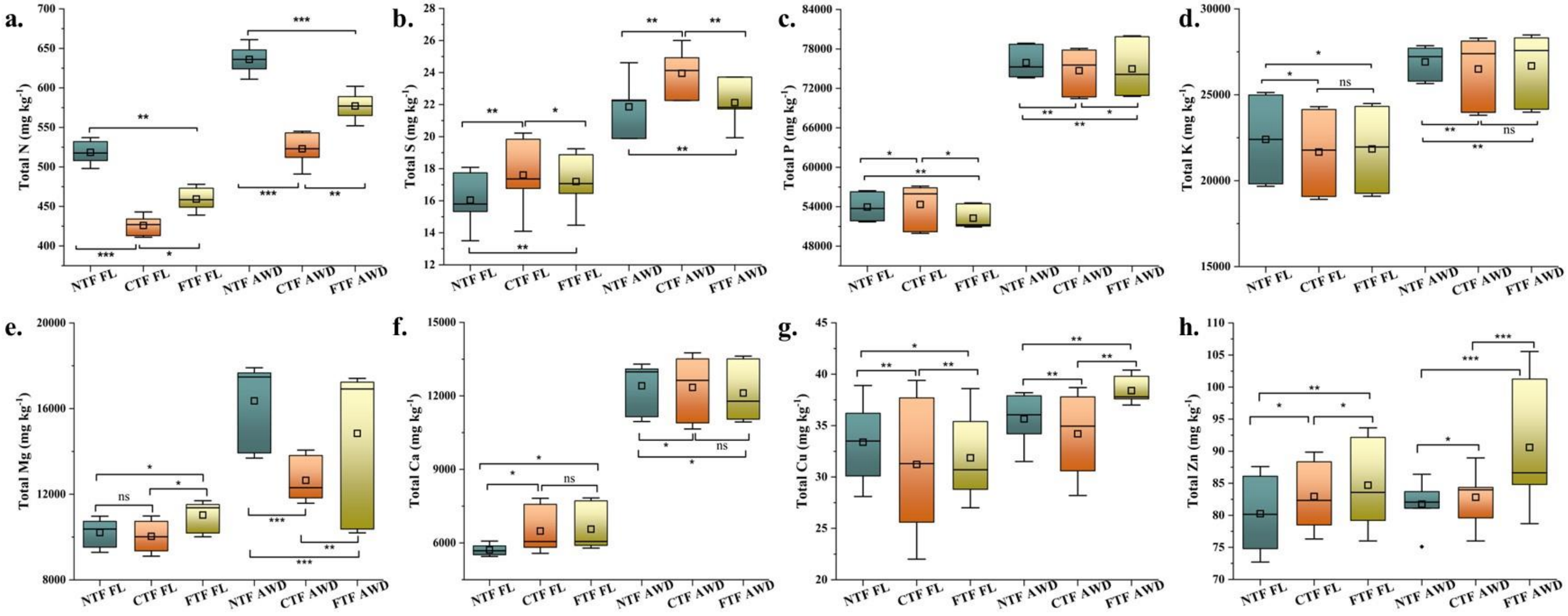

**Extended figure 10.** Total soil elemental concentrations analysed from three field setups with differential irrigation.

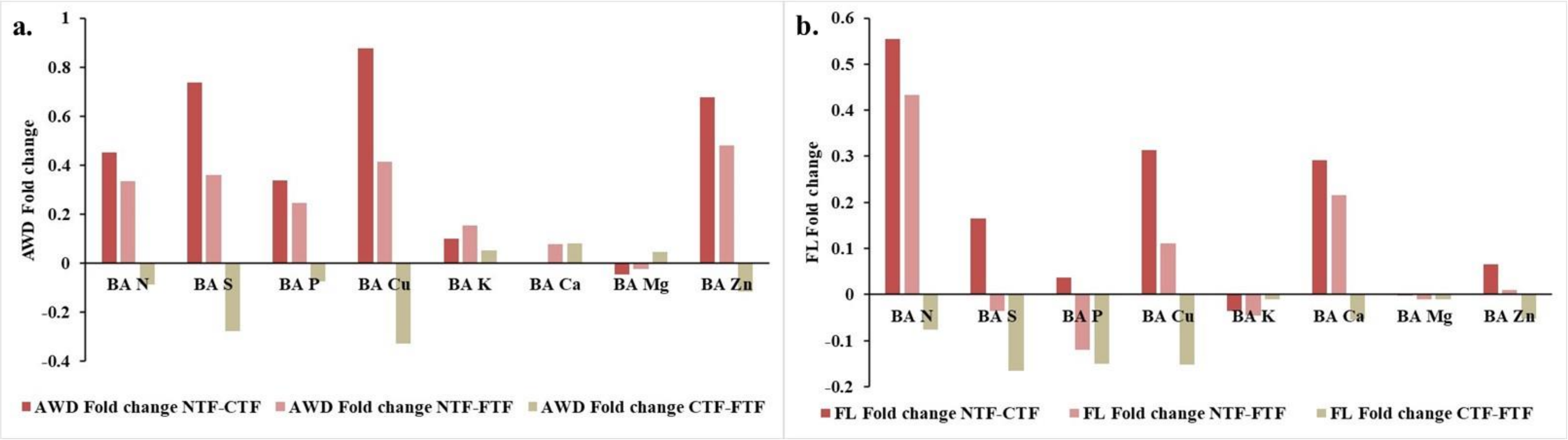

**Extended figure 11.** Fold change of the bioavailability in three field setups for the selected elements under AWD and FL irrigations.

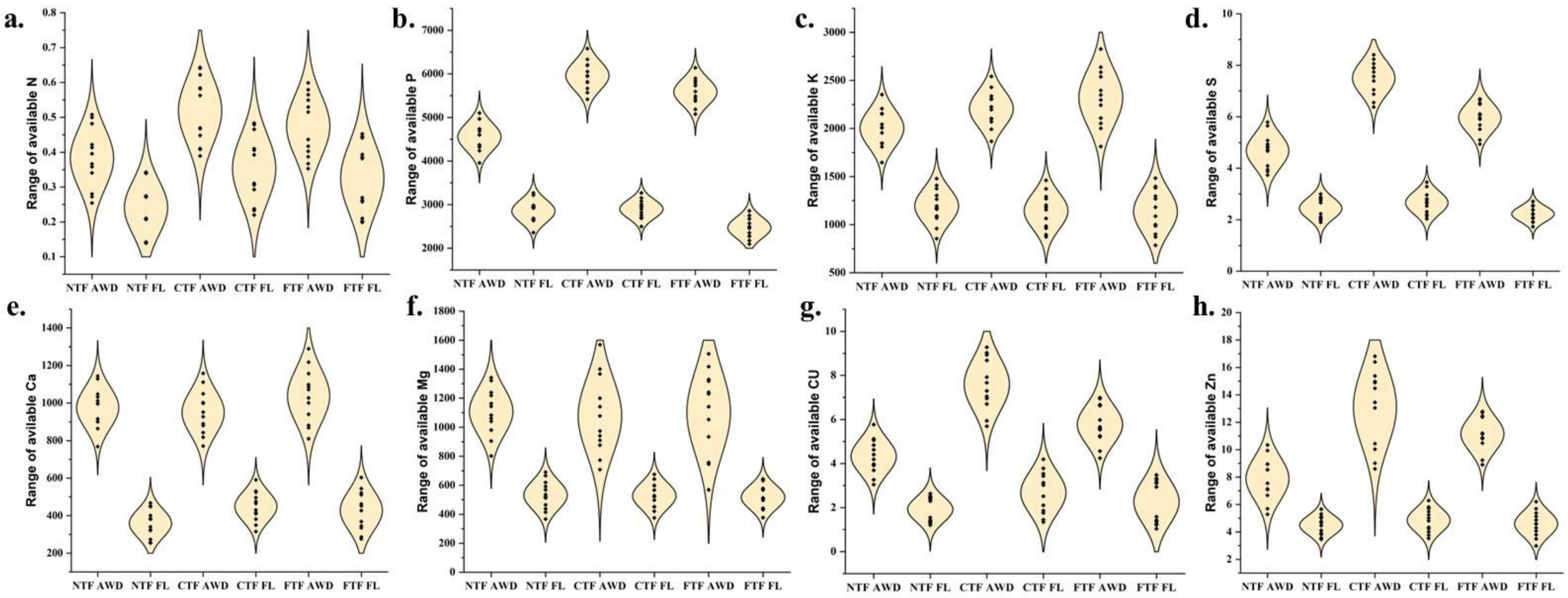

**Extended figure 12**. Average data trend of bioavailable elements in the normal distribution model.

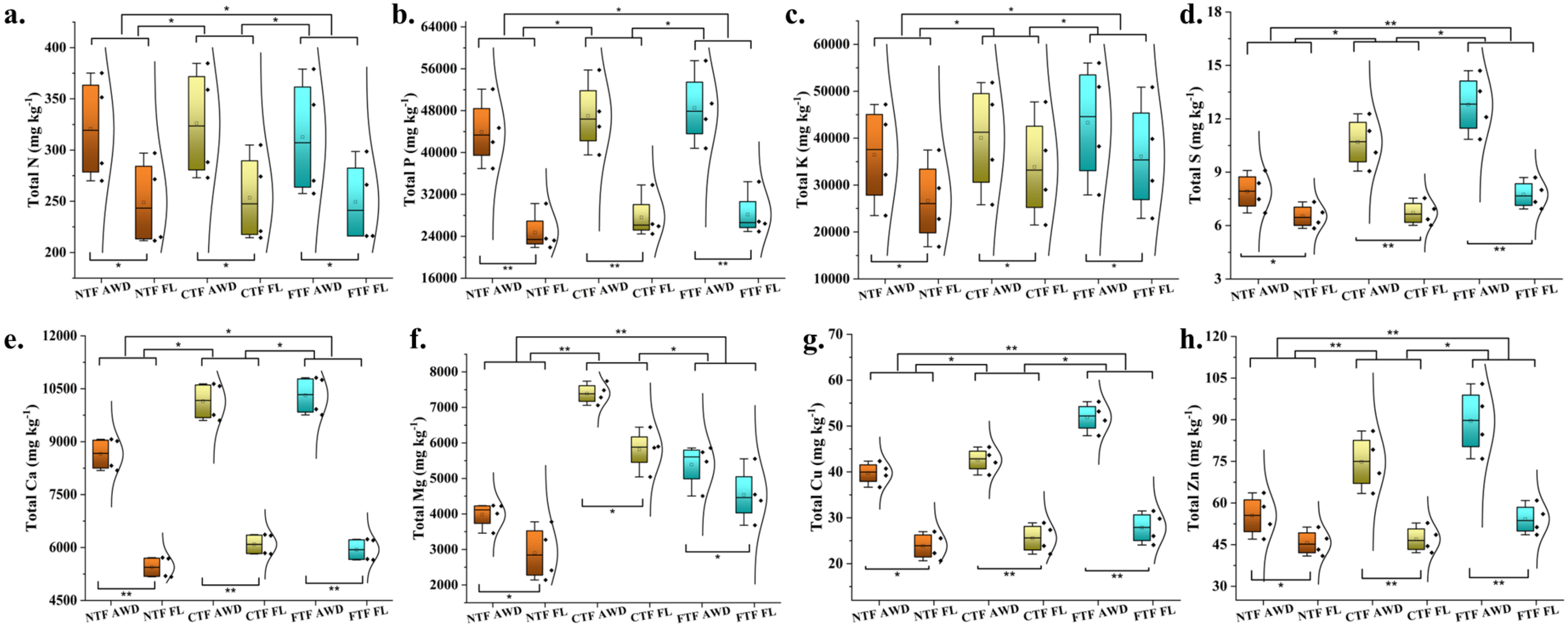

**Extended figure 13.** Total plant elemental concentrations analysed from three field setups with differential irrigation at the final harvest phase.

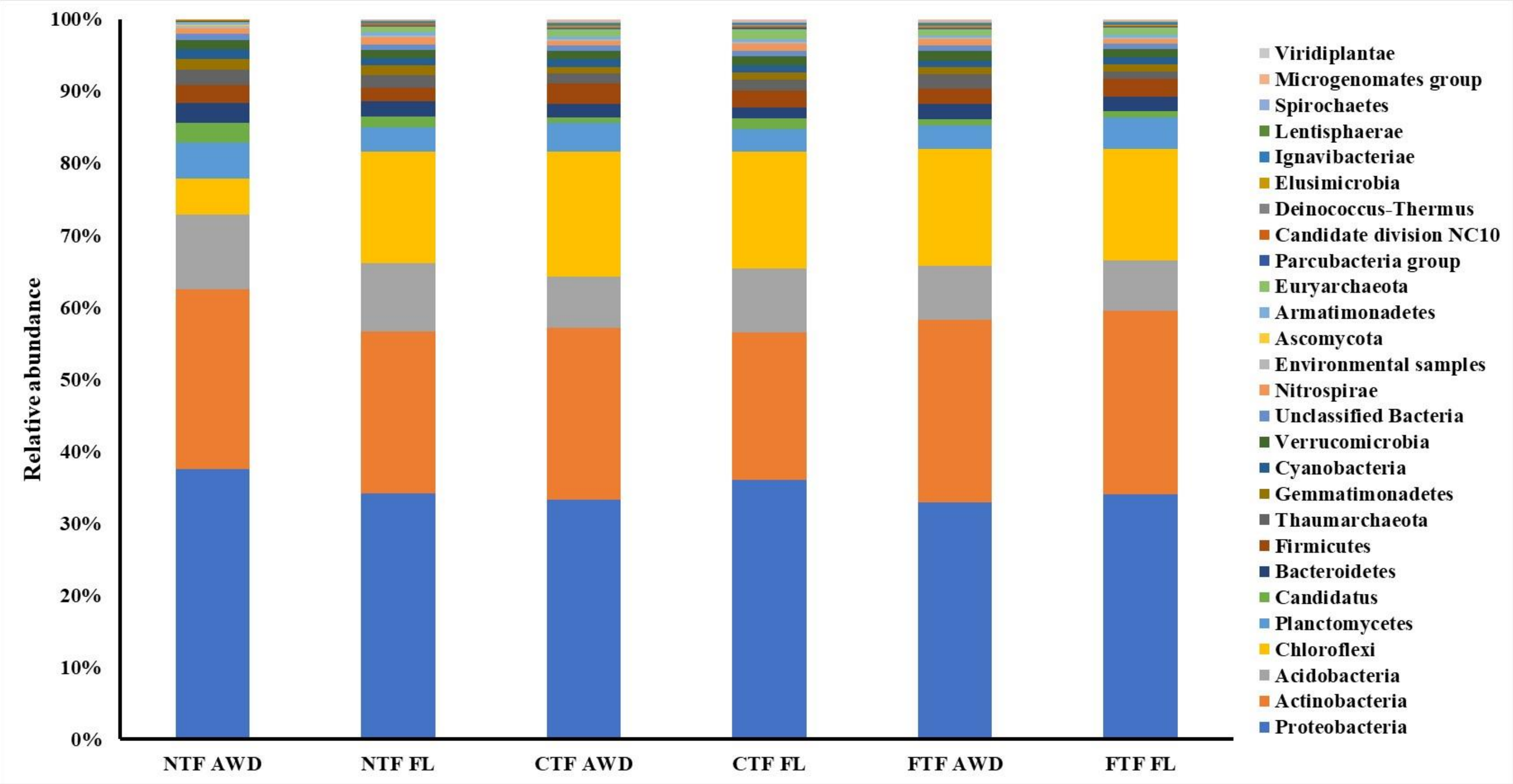

**Extended figure 14.** Microbial phyla relative abundance from the read-counts presented in stacked columns with a color-coded formation.

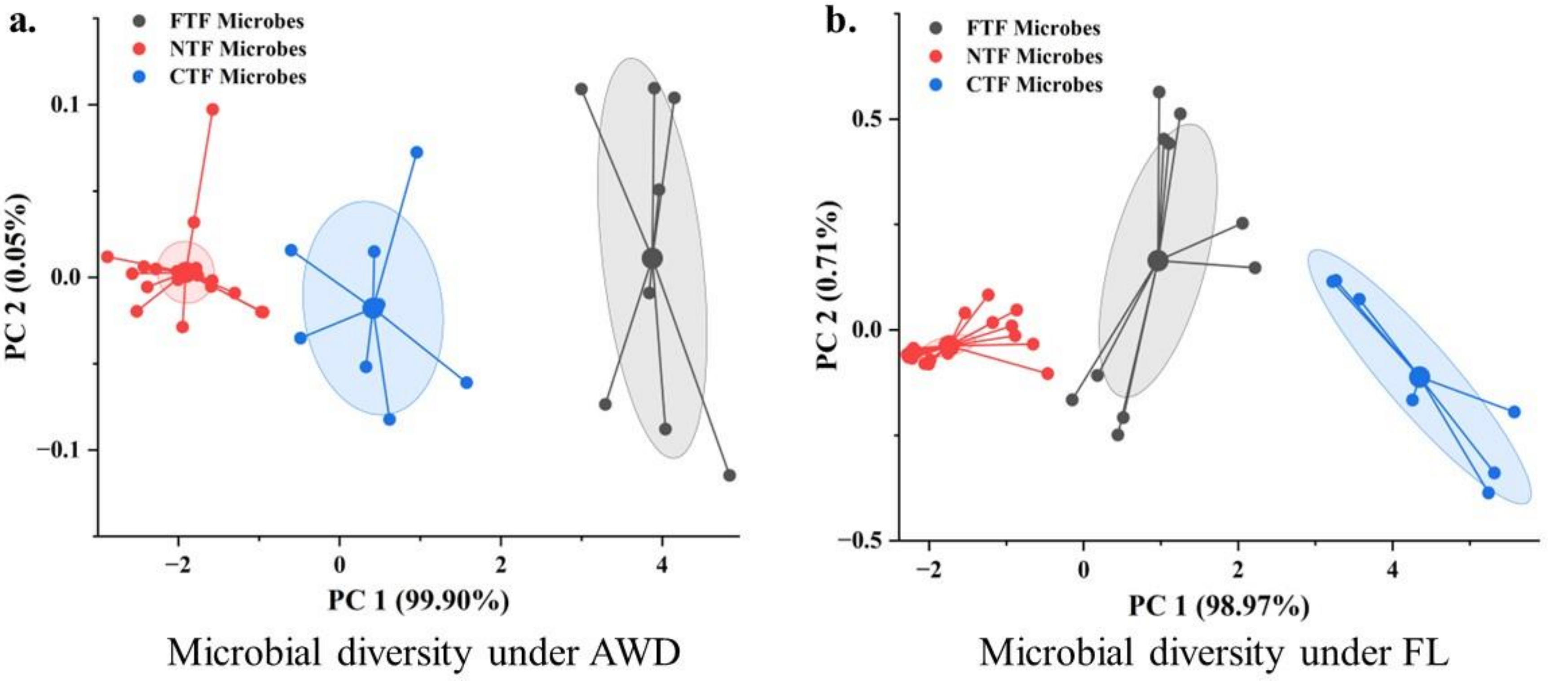

**Extended figure 15**. Machine-learning-based K-means cluster analysis for the statistical variance and significance analysis combined with principal component (PC) analysis.

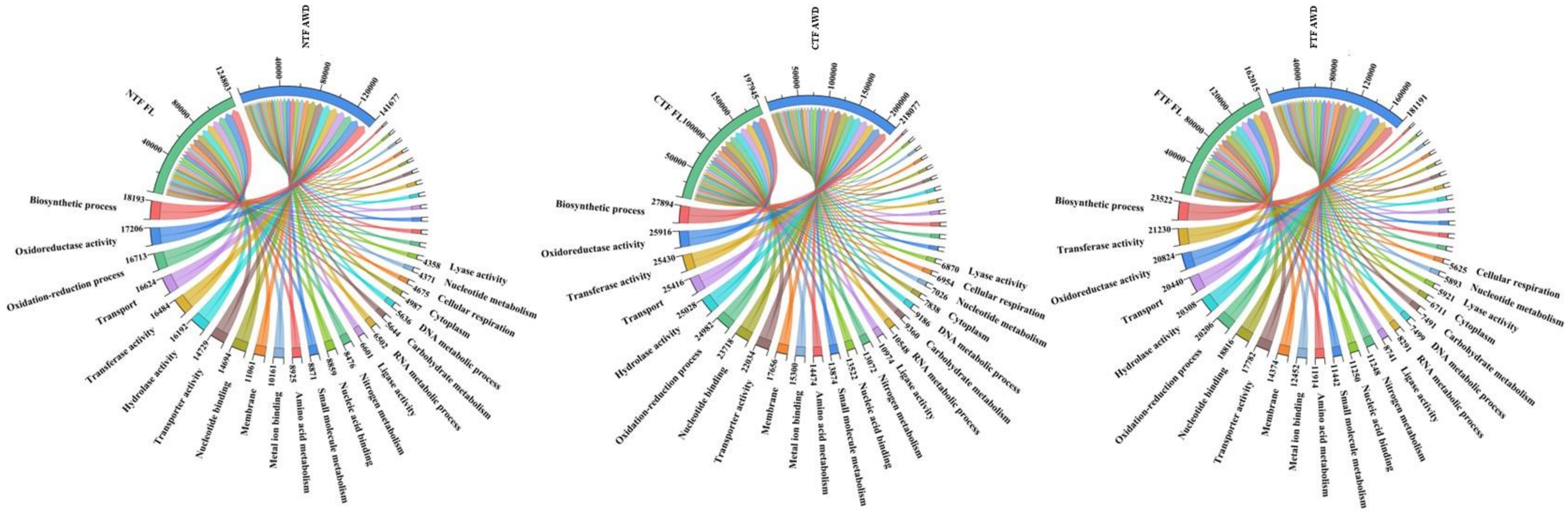

**Extended figure 16.** High frequency gene ontological terms found in microbial communities from the three field setups with varied irrigation.

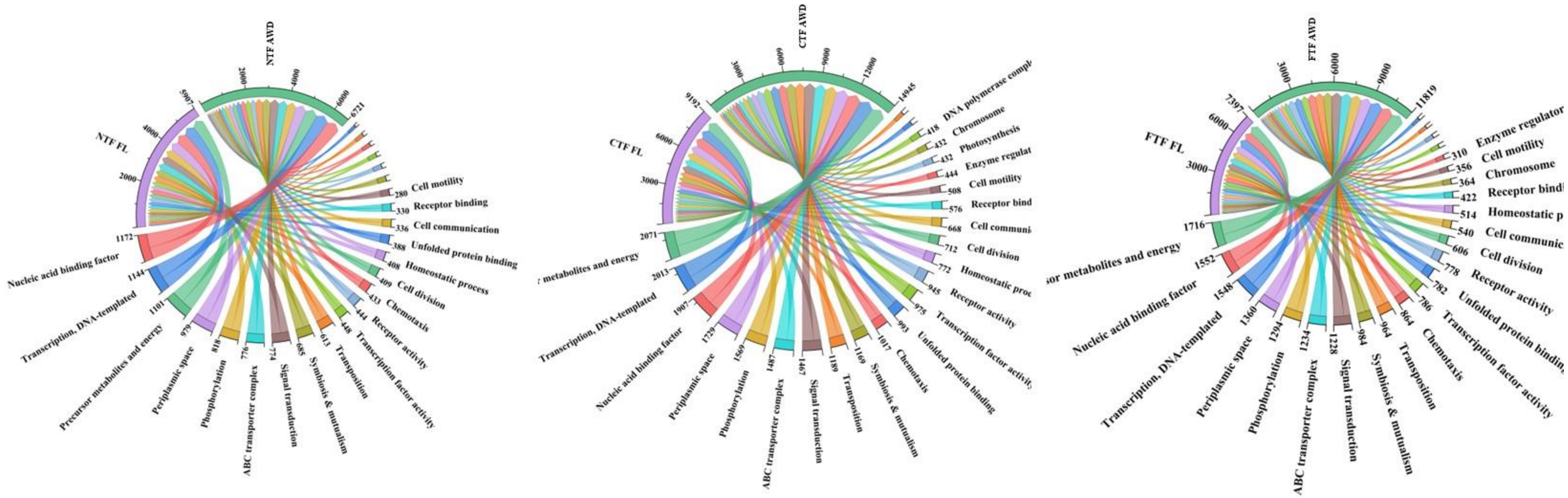

**Extended figure 17.** Low frequency gene ontological terms found in microbial communities from the three field setups with varied irrigation.

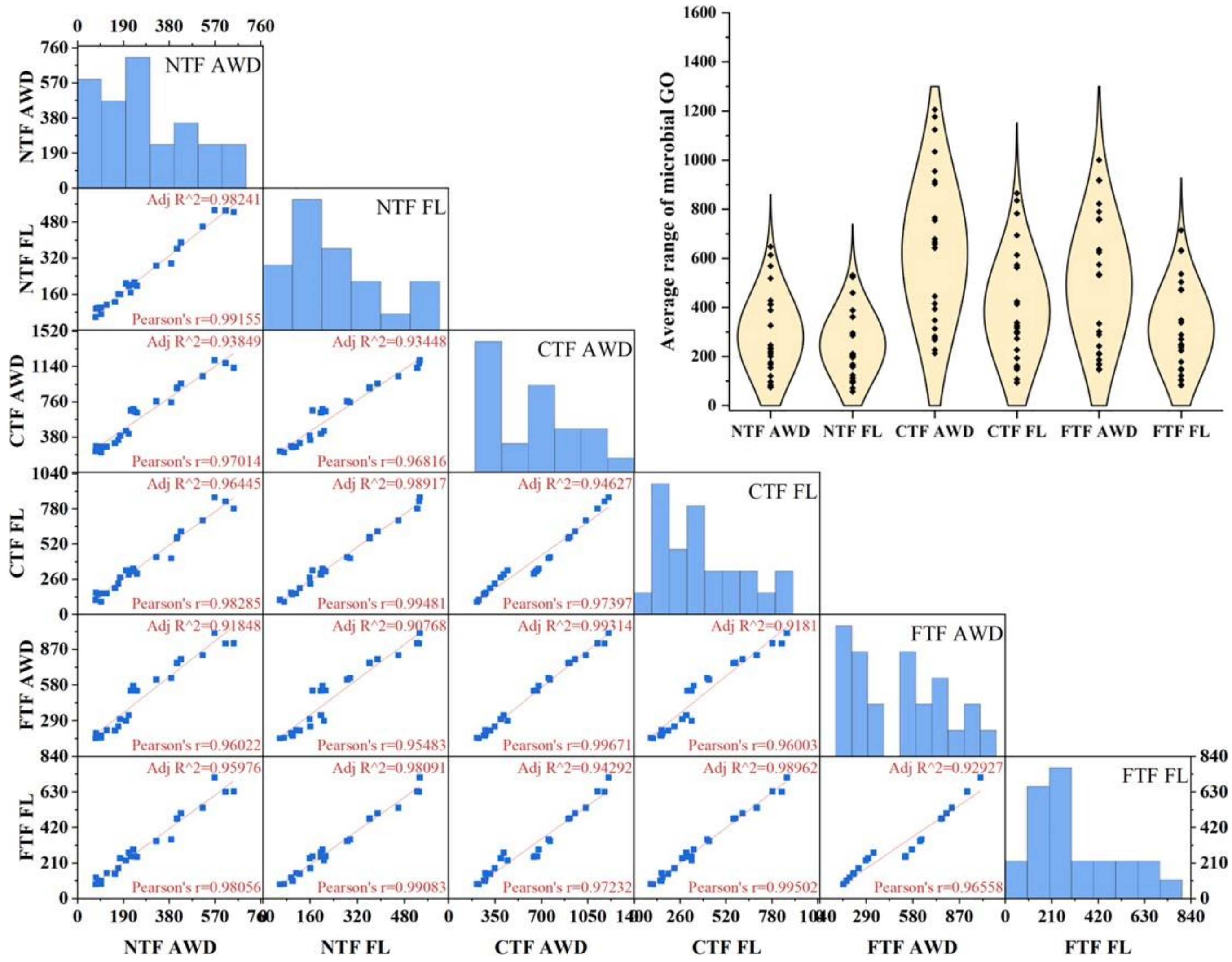

**Extended figure 18**. Statistical analysis of microbial GO distribution in differential setups with a multi-level scatter-matrix plot with linear fit. The violin plot shows the normal distribution mode of the average GO terms counted for these setups.

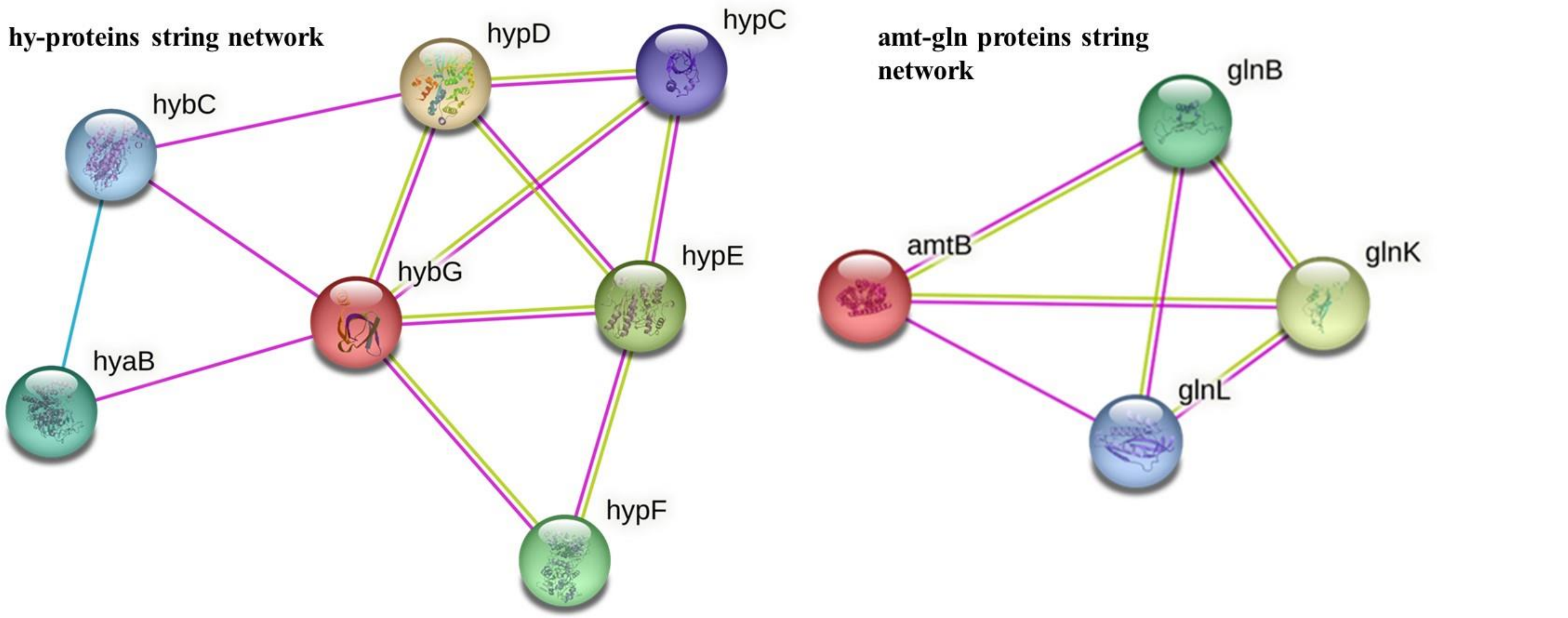

**Extended figure 19.** String networks of hyb-hyp proteins and amt-gln proteins that participate in different metal and gaseous molecule binding and transportation at the cellular level.

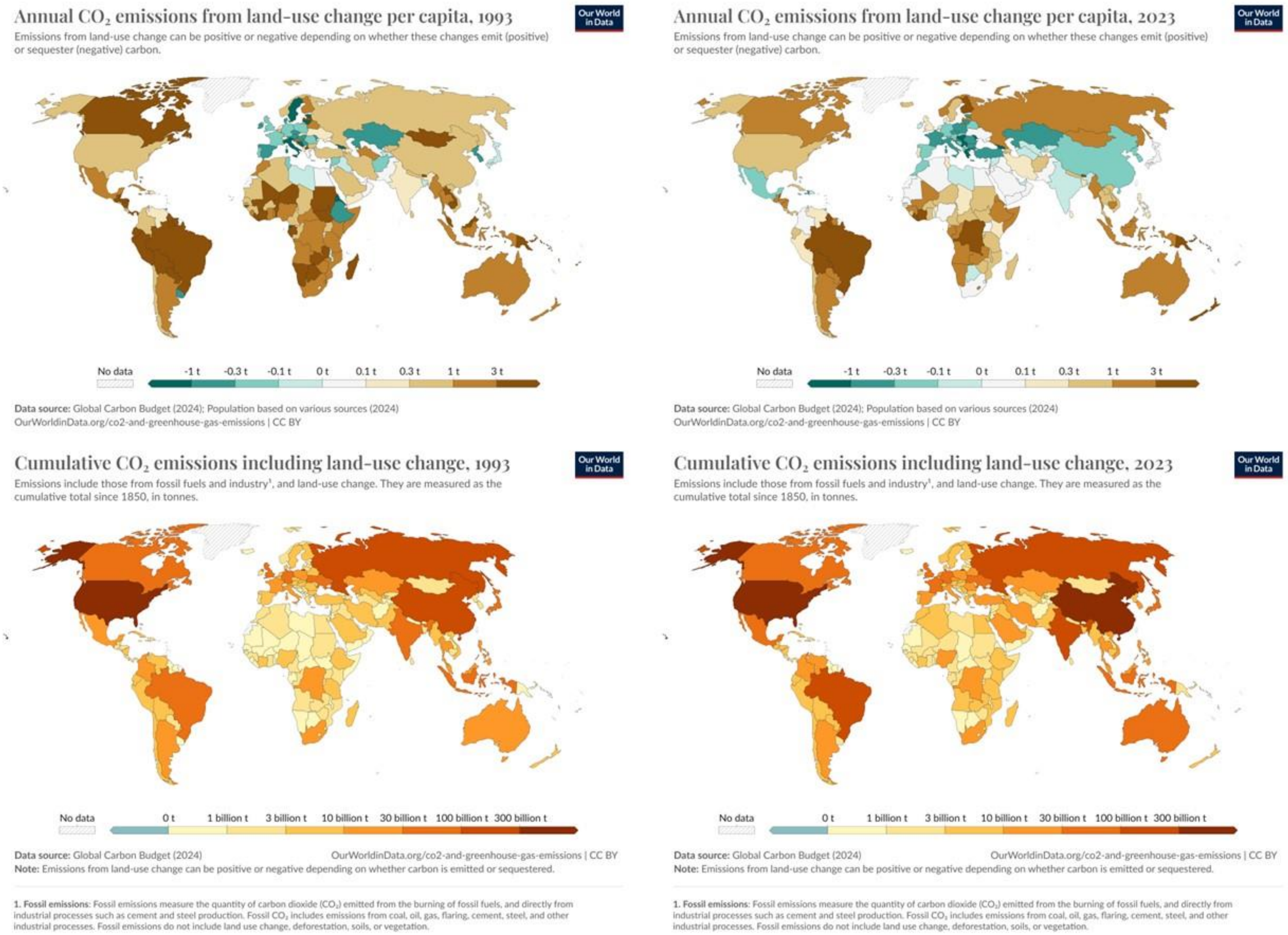

**Extended figure 20.** Annual and cumulative $CO_2$ release global map within the span of 30 years (1993-2023).

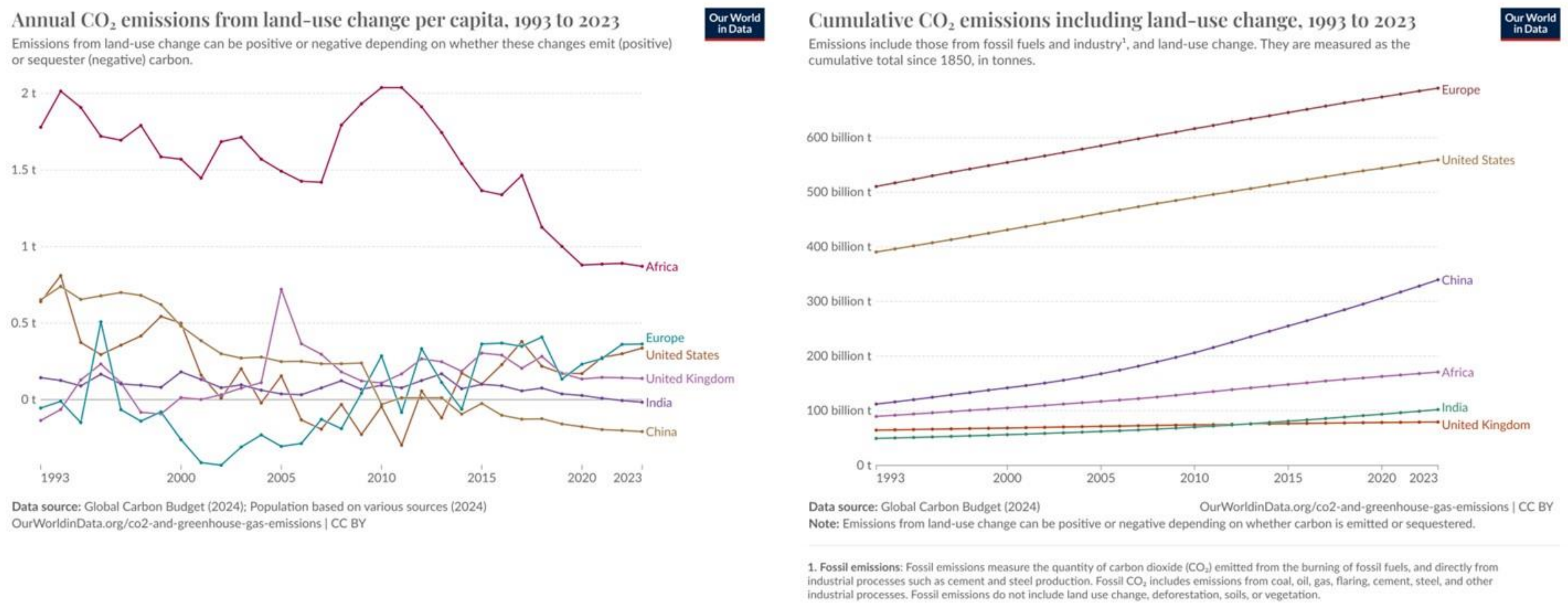

**Extended figure 21.** Annual and cumulative $CO_2$ release trend chart within the span of 30 years (1993-2023).

| $CO_2$ collection and field trial sites | | | | | | | |
|---|---|---|---|---|---|---|---|
| | | | Coordinates | | | | Coordinates |
| | | | Latitude & Longitude | | | | Latitude & Longitude |
| 2022 | Area 1 | AWD 1st | 22.966965, 88.588591 | 2023 | Area 1 | AWD 1st | 22.966965, 88.588591 |
| | | AWD 2nd | 22.965809, 88.588559 | | | AWD 2nd | 22.965809, 88.588559 |
| | | AWD 3rd | 22.966194, 88.590125 | | | AWD 3rd | 22.966194, 88.590125 |
| | | FL 1st | 22.966925, 88.587014 | | | FL 1st | 22.966925, 88.587014 |
| | | Fl 2nd | 22.966422, 88.584933 | | | Fl 2nd | 22.966422, 88.584933 |
| | | Fl 3rd | 22.965295, 88.586714 | | | Fl 3rd | 22.965295, 88.586714 |
| 2022 | Area 2 | AWD 1st | 22.971983, 88.586649 | 2023 | Area 2 | AWD 1st | 22.971983, 88.586649 |
| | | AWD 2nd | 22.971716, 88.588817 | | | AWD 2nd | 22.971716, 88.588817 |
| | | AWD 3rd | 22.970155, 88.586317 | | | AWD 3rd | 22.970155, 88.586317 |
| | | FL 1st | 22.974077, 88.582755 | | | FL 1st | 22.974077, 88.582755 |
| | | Fl 2nd | 22.974255, 88.584321 | | | Fl 2nd | 22.974255, 88.584321 |
| | | Fl 3rd | 22.972348, 88.584032 | | | Fl 3rd | 22.972348, 88.584032 |
| 2022 | Area 3 | AWD 1st | 22.962342, 88.579354 | 2023 | Area 3 | AWD 1st | 22.962342, 88.579354 |
| | | AWD 2nd | 22.962105, 88.581371 | | | AWD 2nd | 22.962105, 88.581371 |
| | | AWD 3rd | 22.960563, 88.580169 | | | AWD 3rd | 22.960563, 88.580169 |
| | | FL 1st | 22.962095, 88.576929 | | | FL 1st | 22.962095, 88.576929 |
| | | Fl 2nd | 22.960771, 88.577680 | | | Fl 2nd | 22.960771, 88.577680 |
| | | Fl 3rd | 22.960060, 88.579482 | | | Fl 3rd | 22.960060, 88.579482 |
| 2022 | Area 4 | AWD 1st | 22.963092, 88.587357 | 2023 | Area 4 | AWD 1st | 22.963092, 88.587357 |
| | | AWD 2nd | 22.963616, 88.589160 | | | AWD 2nd | 22.963616, 88.589160 |

| | | | | | | | |
|---|---|---|---|---|---|---|---|
| | | AWD 3rd | 22.961848, 88.588634 | | | AWD 3rd | 22.961848, 88.588634 |
| | | FL 1st | 22.960544, 88.585308 | | | FL 1st | 22.960544, 88.585308 |
| | | Fl 2nd | 22.960988, 88.588430 | | | Fl 2nd | 22.960988, 88.588430 |
| | | Fl 3rd | 22.959329, 88.586692 | | | Fl 3rd | 22.959329, 88.586692 |
| 2022 | Area 5 | AWD 1st | 22.952700, 88.585534 | 2023 | Area 5 | AWD 1st | 22.952700, 88.585534 |
| | | AWD 2nd | 22.953757, 88.587797 | | | AWD 2nd | 22.953757, 88.587797 |
| | | AWD 3rd | 22.952502, 88.586907 | | | AWD 3rd | 22.952502, 88.586907 |
| | | FL 1st | 22.953895, 88.580781 | | | FL 1st | 22.953895, 88.580781 |
| | | Fl 2nd | 22.953885, 88.583109 | | | Fl 2nd | 22.953885, 88.583109 |
| | | Fl 3rd | 22.952650, 88.581371 | | | Fl 3rd | 22.952650, 88.581371 |
| 2022 | Area 6 | AWD 1st | 22.966984, 88.602335 | 2023 | Area 6 | AWD 1st | 22.966984, 88.602335 |
| | | AWD 2nd | 22.967024, 88.604384 | | | AWD 2nd | 22.967024, 88.604384 |
| | | AWD 3rd | 22.965858, 88.604363 | | | AWD 3rd | 22.965858, 88.604363 |
| | | FL 1st | 22.966154, 88.599513 | | | FL 1st | 22.966154, 88.599513 |
| | | Fl 2nd | 22.965957, 88.601445 | | | Fl 2nd | 22.965957, 88.601445 |
| | | Fl 3rd | 22.964742, 88.600243 | | | Fl 3rd | 22.964742, 88.600243 |
| 2022 | Area 7 | AWD 1st | 22.983115, 88.578635 | 2023 | Area 7 | AWD 1st | 22.983115, 88.578635 |
| | | AWD 2nd | 22.983085, 88.580159 | | | AWD 2nd | 22.983085, 88.580159 |
| | | AWD 3rd | 22.981337, 88.579601 | | | AWD 3rd | 22.981337, 88.579601 |
| | | FL 1st | 22.983757, 88.581929 | | | FL 1st | 22.983757, 88.581929 |
| | | Fl 2nd | 22.983549, 88.584128 | | | Fl 2nd | 22.983549, 88.584128 |
| | | Fl 3rd | 22.981919, 88.583131 | | | Fl 3rd | 22.981919, 88.583131 |
| 2022 | Area 8 | AWD 1st | 22.988912, 88.613879 | 2023 | Area 8 | AWD 1st | 22.988912, 88.613879 |

| | | | | | | | |
|---|---|---|---|---|---|---|---|
| | | AWD 2nd | 22.988586, 88.616594 | | | AWD 2nd | 22.988586, 88.616594 |
| | | AWD 3rd | 22.987520, 88.615113 | | | AWD 3rd | 22.987520, 88.615113 |
| | | FL 1st | 22.986206, 88.612828 | | | FL 1st | 22.986206, 88.612828 |
| | | Fl 2nd | 22.985673, 88.615811 | | | Fl 2nd | 22.985673, 88.615811 |
| | | Fl 3rd | 22.984359, 88.616411 | | | Fl 3rd | 22.984359, 88.616411 |
| 2022 | Area 9 | AWD 1st | 22.994058, 88.614673 | 2023 | Area 9 | AWD 1st | 22.994058, 88.614673 |
| | | AWD 2nd | 22.994364, 88.617034 | | | AWD 2nd | 22.994364, 88.617034 |
| | | AWD 3rd | 22.992656, 88.616519 | | | AWD 3rd | 22.992656, 88.616519 |
| | | FL 1st | 22.994572, 88.619673 | | | FL 1st | 22.994572, 88.619673 |
| | | Fl 2nd | 22.994483, 88.621744 | | | Fl 2nd | 22.994483, 88.621744 |
| | | Fl 3rd | 22.993456, 88.620113 | | | Fl 3rd | 22.993456, 88.620113 |
| 2022 | Area 10 | AWD 1st | 22.970590, 88.603998 | 2023 | Area 10 | AWD 1st | 22.970590, 88.603998 |
| | | AWD 2nd | 22.970135, 88.606004 | | | AWD 2nd | 22.970135, 88.606004 |
| | | AWD 3rd | 22.968683, 88.604792 | | | AWD 3rd | 22.968683, 88.604792 |
| | | FL 1st | 22.970777, 88.607796 | | | FL 1st | 22.970777, 88.607796 |
| | | Fl 2nd | 22.970234, 88.610060 | | | Fl 2nd | 22.970234, 88.610060 |
| | | Fl 3rd | 22.969009, 88.608311 | | | Fl 3rd | 22.969009, 88.608311 |
| 2022 | Area 11 | AWD 1st | 22.952179, 88.569017 | 2023 | Area 11 | AWD 1st | 22.952179, 88.569017 |
| | | AWD 2nd | 22.952742, 88.571431 | | | AWD 2nd | 22.952742, 88.571431 |
| | | AWD 3rd | 22.951033, 88.570744 | | | AWD 3rd | 22.951033, 88.570744 |
| | | FL 1st | 22.951428, 88.566527 | | | FL 1st | 22.951428, 88.566527 |
| | | Fl 2nd | 22.951340, 88.564864 | | | Fl 2nd | 22.951340, 88.564864 |
| | | Fl 3rd | 22.949917, 88.566045 | | | Fl 3rd | 22.949917, 88.566045 |

| | | | | | | | |
|---|---|---|---|---|---|---|---|
| 2022 | Area 12 | AWD 1st | 22.955706, 88.582996 | 2023 | Area 12 | AWD 1st | 22.955706, 88.582996 |
| | | AWD 2nd | 22.955242, 88.585260 | | | AWD 2nd | 22.955242, 88.585260 |
| | | AWD 3rd | 22.953632, 88.582664 | | | AWD 3rd | 22.953632, 88.582664 |
| | | FL 1st | 22.955183, 88.578705 | | | FL 1st | 22.955183, 88.578705 |
| | | Fl 2nd | 22.955173, 88.580314 | | | Fl 2nd | 22.955173, 88.580314 |
| | | Fl 3rd | 22.953661, 88.580003 | | | Fl 3rd | 22.953661, 88.580003 |

**Extended table 1**. Coordinates of the selected 12 experimental sites where field trials were conducted in two consecutive years. Two irrigation regimes, alternate wetting and drying (AWD) and flooded (FL), were combined with three tillage practices.

| Soil $CO_2$ | | |
|---|---|---|
| Mann-Whitney U Test at $P<0.05$ | | |
| | U values | Exact *P* values |
| NTF AWD to CTF AWD | 19 | 0.00382 |
| NTF FL to CTF FL | 30 | 0.00414 |
| NTF AWD to FTF AWD | 77 | 0.0069 |
| NTF FL to FTF FL | 72 | 0.00732 |
| CTF AWD to FTF AWD | 32 | 0.08242 |
| CTF FL to FTF FL | 41 | 0.07928 |

**Extended table 2**. Post-hoc analysis of soil $CO_2$ release from the fields with different combinations.

| Soil elements | | |
| --- | --- | --- |
| Mann-Whitney U Test at *P*<0.05 | | |
| | U values | Exact *P* values |
| NTF AWD to CTF AWD | 29 | 0.00472 |
| NTF FL to CTF FL | 34 | 0.00428 |
| NTF AWD to FTF AWD | 81 | 0.00688 |
| NTF FL to FTF FL | 56 | 0.00722 |
| CTF AWD to FTF AWD | 52 | 0.00826 |
| CTF FL to FTF FL | 61 | 0.00794 |

**Extended table 3**. Post-hoc analysis of soil elemental bioavailability from the fields with different combinations.

| Soil Microbes | | |
| --- | --- | --- |
| Mann-Whitney U Test at *P*<0.05 | | |
| | U values | Exact *P* values |
| NTF AWD to CTF AWD | 22 | 0.00212 |
| NTF FL to CTF FL | 30 | 0.00218 |
| NTF AWD to FTF AWD | 71 | 0.00422 |
| NTF FL to FTF FL | 42 | 0.00436 |
| CTF AWD to FTF AWD | 49 | 0.00596 |
| CTF FL to FTF FL | 53 | 0.00608 |

**Extended table 4**. Post-hoc analysis of soil microbial diversity changes in the fields with different combinations.

| Years | Tillage practice | Variogram model | Coefficient of determination (%) | Lag distance (m) | Maximum distance (m) |
|---|---|---|---|---|---|
| 2022 | NTF | -0.0231004 + 0.00699175 ln(1 + x) | 53.90 % | 171.50 | 3822.68 |
| | CTF | -0.171484 + 0.0439949 ln(1 + x) | 66.33 % | 171.50 | 3822.68 |
| | FTF | -0.0911852 + 0.0186986 ln(1 + x) | 81.74% | 171.50 | 3822.68 |
| 2023 | NTF | -0.0466559 + 0.0151919 ln(1 + x) | 35.54% | 171.50 | 3822.68 |
| | CTF | -0.288862 + 0.0823526 ln(1 + x) | 56.32% | 171.50 | 3822.68 |
| | FTF | -0.152707 + 0.0363219 ln(1 + x) | 58.41% | 171.50 | 3822.68 |

**Extended table 5**. Variogram models for Ordinary Kriging corresponding to various tillage practices.
